# Pre-division TCF1 drop determines long-term CD8 T cell fates

**DOI:** 10.1101/2025.10.06.680655

**Authors:** Natalie R. Favret, Maider Astorkia, Melissa M. Wolf, Michael W. Rudloff, Claudia N. McDavid, Megan M. Erwin, Kristen A. Murray, Jessica J. Roetman, Carlos R. Detrés Román, Zachary D. Ewell, Lincoln A. Brown, William T. Murray, Paul Zumbo, Doron Betel, Mary Philip

## Abstract

T Cell Factor 1 (TCF1) is a master transcription factor controlling T cell development and peripheral T cell differentiation during infection, cancer, and autoimmunity. TCF1 is highly expressed in naive CD8 T cells but must be downregulated as T cells proliferate to become effectors. If and how TCF1 plays a role during T cell priming prior to cell cycle entry is unknown. Surprisingly, we found that TCF1 expression is rapidly downregulated within hours after antigen encounter in both murine and human CD8 T cells, even before T cells enter cell cycle. TCF1 then rebounds to high levels upon cell cycle entry, ultimately declining again with proliferation and effector differentiation. This rapid pre-division drop and rebound occurs in diverse settings, including infection and cancer. The magnitude of the pre-division TCF1 drop is modulated by TCR signal strength and inflammatory cytokines and strikingly, regulates long-term effector and memory fates. Paired transcriptomic and epigenetic analyses revealed that TCF1-regulated chromatin regions were remodeled within hours following antigen encounter, activating effector and inflammatory cytokine signaling modules and poising T cells for effector differentiation. Remarkably, transient siRNA-mediated TCF1 downmodulation during the pre-division priming phase was sufficient to induce long-term population effector skewing. We have uncovered a novel mechanism whereby pre-division dynamic TCF1 regulation determines long-term CD8 T cell fate commitment, potentially serving as a critical checkpoint regulating T cell responses in infection, cancer, and autoimmunity.

## INTRODUCTION

Naive CD8 T cells differentiating in response to acute infection must balance divergent cell fates, giving rise to many short-lived cytotoxic effectors and a small number of long-lived memory T cells. Failure to generate effectors limits pathogen clearance, whereas inadequate memory formation compromises long-term immunity. The naive CD8 T cell pool is highly plastic, with single antigen-specific T cells able to give rise to heterogeneous populations of effector and memory cells^1,2^. There have been many studies and much debate on the regulation of T cell differentiation to effector and memory fates and the lineage relationship between naive, effector, and memory T cells^3^. A shared feature of the current models is that T cell differentiation occurs as T cells proliferate. Even the earliest fate decision models such as the asymmetric division model invoke the process of mitosis/cytokinesis as the origin of different fates^4,5^.

We recently showed that CD8 T cell selection between the functional effector fate in acute infection versus the dysfunctional fate in tumors occurs within hours of activation and before T cells divide^6^. To understand how T cells decide between divergent fates, we set out to determine the earliest molecular programs engaged when T cells are activated and before they enter cell cycle. Transcription factors (TF) are key mediators of cell fate decisions in embryonic and adult somatic cell differentiation^7^, including T cells^8^. We focused on T cell factor 1 (TCF1), a master regulator of T cell fate^9,10^ expressed at high levels in naive T cells^11^. TCF1 is known to be downregulated as T cells proliferate and differentiate into functional effectors^12-16^, however, its regulation during the early priming phase prior to cell cycle entry has not been studied.

Here, we examined the expression of *Tcf7* mRNA (encoding TCF1) and TCF1 protein in the earliest hours after priming of antigen-specific CD8 T cells during acute infection or in tumors. Unexpectedly, *Tcf7* mRNA/TCF1 fell sharply within hours of activation and even prior to cell division, only to rebound again as T cells entered cell cycle. We dissected the regulation of this rapid, pre-division TCF1 drop and sought to determine its impact on long-term T cell differentiation. We found that TCR signaling strength and inflammatory cytokines modulate the pre-division TCF1 drop, determining long-term population skewing between the effector and memory fates. The transient pre-division TCF1 drop resulted in chromatin remodeling and expression changes in TCF1-regulated genes, including genes required for T cell effector differentiation. TCF1-mediated pre-division molecular programming was further reinforced by inflammatory cytokines as T cells proliferate, leading to effector fate commitment. Thus, the pre-division priming-induced TCF1 drop represents a distinct and previously uncharacterized first regulatory step on the road to T cell fate commitment.

## RESULTS

### TCF1 drops upon CD8 T cell priming prior to cell division

To determine how *Tcf7* mRNA is regulated during priming, we analyzed our previously-published RNA-SEQ data obtained from antigen-specific CD8 T cells activated during acute infection with LM^6^. SV40 large T antigen (TAG) epitope-I (TAG_206-215_)-specific CD8 T cells (TCR_TAG_)^17^ were adoptively transferred into C57BL/6 mice (B6) infected with a TAG-expressing *Listeria monocytogenes* (LM_TAG_) and reisolated and analyzed 6, 12, and 24 hours (h) later (**Fig. 1A**). Unexpectedly, *Tcf7* mRNA was rapidly downregulated in activated CD8 T cells (**Fig. 1B, left**). We confirmed at the protein level that in activated CD69+ cells (**Extended Data** Fig. 1A), TCF1 dropped within 6h and remained low over the next 12h (**Fig. 1B, right**) both in TCR_TAG_ responding to LM_TAG_ acute infection and in (OVA)-specific transgenic CD8 T cells (TCR_OTI_) activated in LM_OVA_-infected B6 (**Extended Data** Fig. 1B). To test whether this early TCF1 drop was specific to the infection context, we analyzed RNA-SEQ data from TCR_TAG_ early after transfer into mice with late-stage TAG-expressing liver tumors^6^. *Tcf7* RNA/TCF1 protein dropped to low levels within 6h of activation in liver tumors (**Extended Data** Fig. 1C), suggesting that this rapid activation-induced TCF1 drop is not context-dependent but in response to cognate antigen encounter. It is well-established that activated CD8 T cells downregulate TCF1 after they have divided 3+ times^18,19^, however few studies have examined pre-division time points *in vivo*.

**Figure 1.**
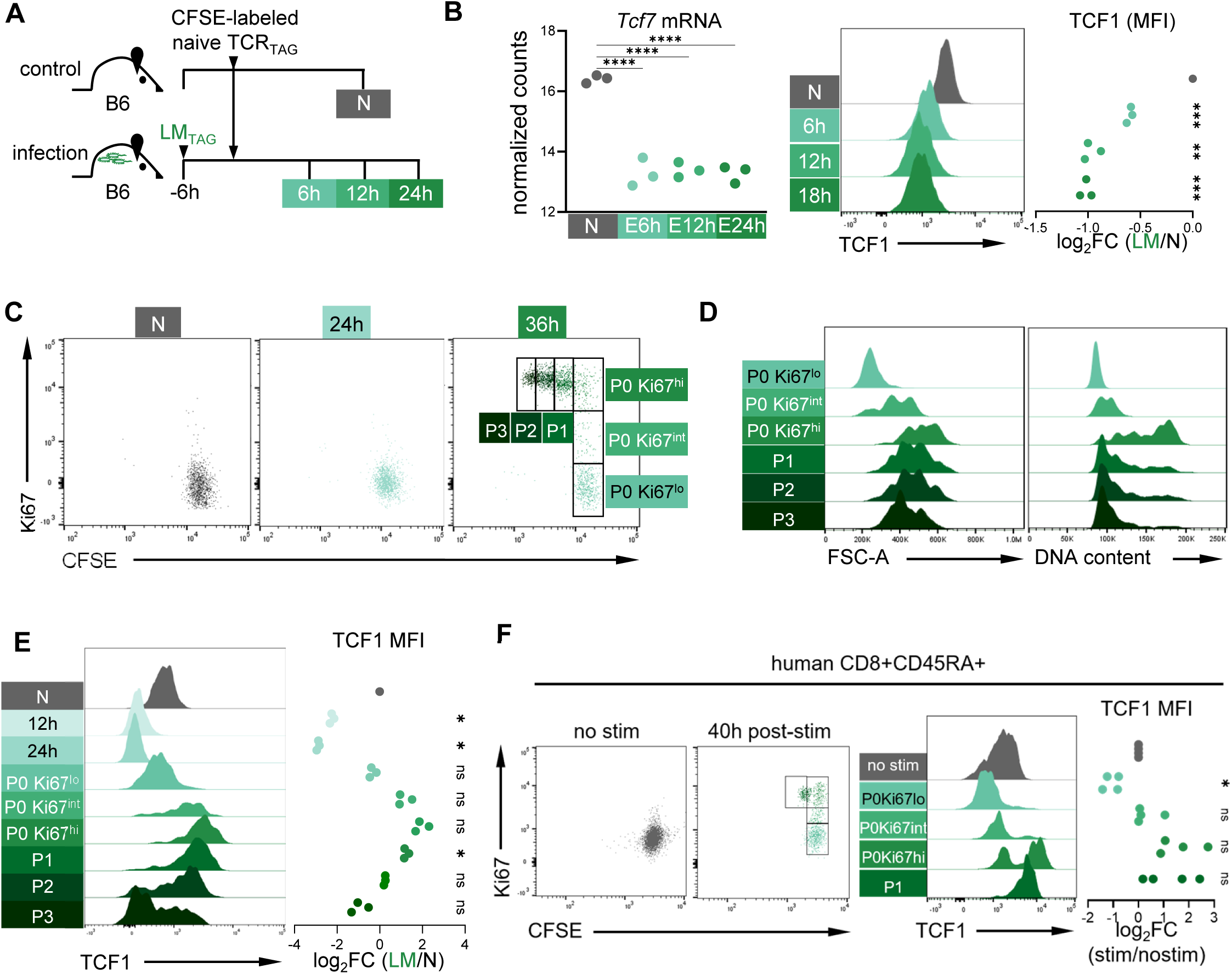
TCF1 drops upon CD8 T cell priming prior to cell division. **A**. Experimental scheme: CFSE-labeled naive TCR_TAG_ (Thy1.1) were adoptively transferred into B6 or LM_TAG_-infected B6 (Thy1.2). Lymphocytes were re-isolated from spleens and analyzed by flow cytometry at 6, 12, and 24 hours (h) post-transfer. **B**. Normalized counts (variance-stabilized (log_2_) expression) of *Tcf7* transcripts from naive TCR_TAG_ and TCR_TAG_ sorted from LM_TAG_ -infected mice 6, 12, and 24h post-transfer from published RNA-SEQ dataset (GSE209712)^12^ (left). Representative histograms of TCF1 expression on CD8^+^ Thy1.1^+^ TCR_TAG_ from spleens of infected mice (green) and naive *in vivo* control (N; grey) (right). Summary plot shows log_2_ fold change (FC) of TCF1 MFI in comparison to N. Each symbol represents an individual mouse with n=3 per group. \*\**P*=0.0024, \*\*\**P*<0.001, \*\*\*\**P*<0.0001 determined by one sample t test. **C**. CFSE and Ki67 expression in CFSE-labeled naive TCR_TAG_ from spleens of infected or naive mice. At 36 hours, cells are gated based on division (P0-P3) and Ki67 expression (lo, int, hi). **D**. FSC-A and DNA content of cell populations gated as in Fig. 1C; in D-E data is concatenated from 3 biologic replicates per condition. **E**. TCF1 expression of cell populations gated as in Fig. 1C. Log_2_ fold change of TCF1 MFI in comparison to N. ns, not significant. \**P*<0.006 determined by one sample t test with Bonferroni correction with n=3 per group. **F.** Human PBMCs were CFSE-labeled and cultured *in vitro* with IL7 (N; grey; no stim) or IL2, aCD3 and aCD28 (green; stim). CFSE, Ki67, and TCF1 expression on CD8^+^CD45RA^+^ after 40h culture. \**P*<0.0125 determined by one sample t test with Bonferroni correction with n=4 per group.

We tracked TCF1 expression over the course of early activation, cell cycle entry, and cell division using CFSE dilution and Ki67^20^, which is rapidly degraded in non-cycling cells^21^. Following transfer into LM_TAG_-infected B6, TCR_TAG_ remained in G0 for at least 24h, and by 36 hours, we could identify undivided cells in G0 (P0 Ki67^low^), G1/S phase (P0 Ki67^int^), and G2/M (P0 Ki67^hi^), as well as Ki67^hi^ cells that had undergone 1-3 divisions (**Fig. 1C**). Increased Ki67 expression in P0 cells correlated with increased cell size (FSC-A) and DNA content (S-phase) (**Fig. 1D**), and early division (P1-P3) remained Ki67^hi^ (**Fig. 1C, D**). TCF1 decreased following activation (12h, 24h) and began rebounding with cell cycle entry (P0 Ki67^int^), with higher-than-naive expression prior to the first division (P0 Ki67^hi^) (**Fig. 1E**). Subsequently, as cells began dividing, TCF1 was again downregulated (**Fig. 1E**). Thus, unexpectedly, TCF1 undergoes *two* rounds of downregulation: a pre-division drop immediately following activation with subsequent rebound on cell cycle entry, and a proliferation-associated secondary drop. To determine whether this pre-division drop and rebound kinetics is conserved in human T cells, we activated CFSE-labeled healthy donor human peripheral blood T cells with anti-CD3 and anti-CD28 monoclonal antibodies and found that TCF1 expression was lowest in undivided Ki67^lo^ CD8 T cells and rebounded as T cells entered cell cycle and upregulated Ki67 (**Fig. 1F**). A recent whole-proteomics analysis also observed a rapid decrease in TCF1 protein within 6h following *in vitro* activation, consistent with our data^22^. We next sought to determine the factors inducing these unexpected dynamic pre-division changes in TCF1 expression.

### Pre-division TCF1 drop is regulated by TCR signaling strength and IL12

Studies have demonstrated that proliferation-associated TCF1 downregulation is controlled by TCR signaling strength and inflammation^19,23,24^. To understand whether the pre-division TCF1 drop is regulated by TCR signal strength, we stimulated TCR_TAG_ *in vitro* using previously characterized TAG_206-215_ altered peptide ligands (APL) with high (wild-type; TAG.N4), intermediate (TAG.F6), and low (TAG.D4) TCR functional avidity^25^. As expected^26^, the proportion of activated CD69+ TCR_TAG_ correlated with TCR functional avidity and peptide concentration (**Fig. 2A left**). Interestingly, the magnitude of the pre-division TCF1 drop in activated TCR_TAG_ correlated with TCR signal strength (**Fig. 2A right**). We confirmed using OVA_257-264_ APL that the pre-division TCF1 drop in TCR_OTI_ similarly increased with greater TCR signal strength (**Fig. 2B right**).

**Figure 2.**
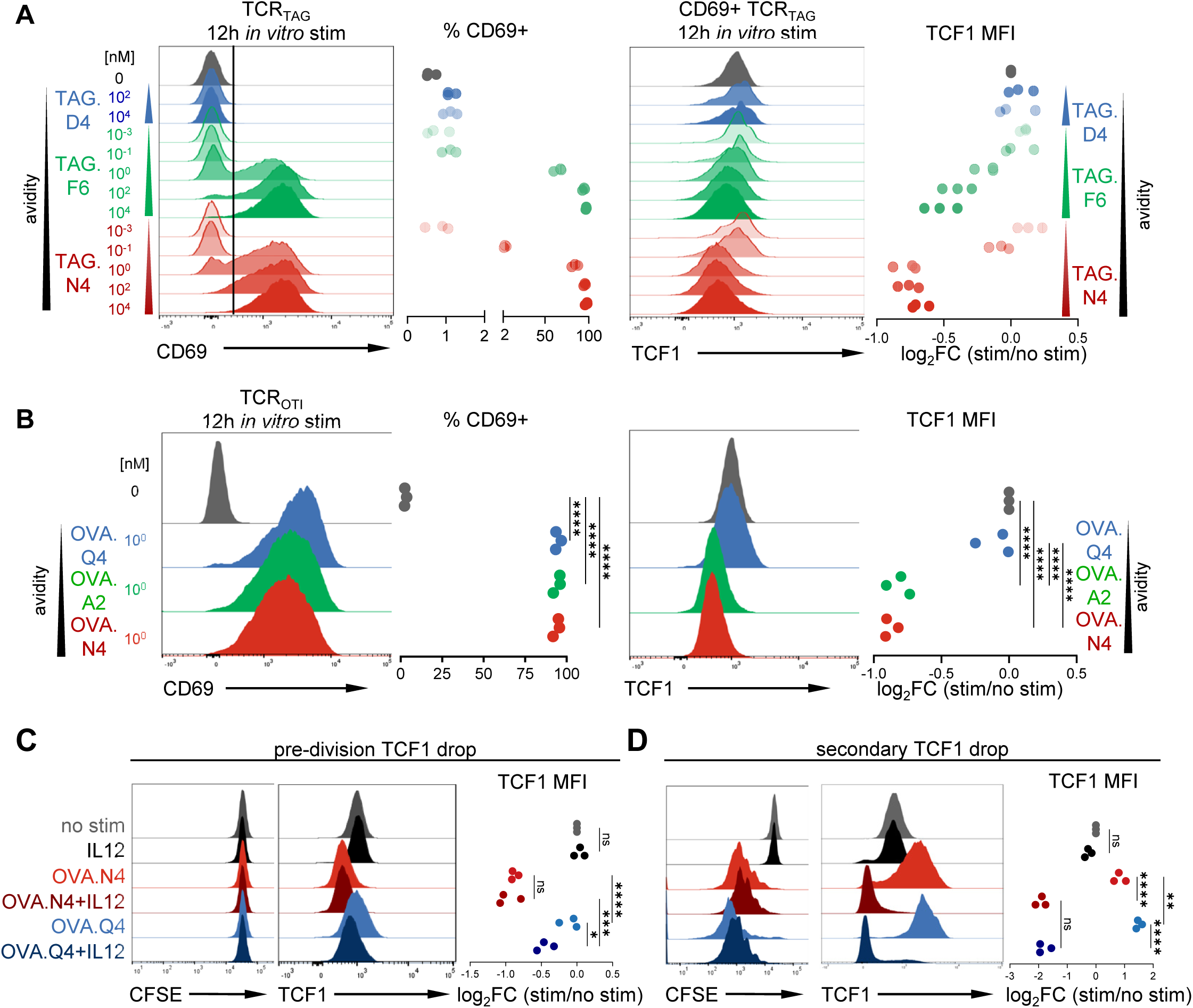
Pre-division TCF1 drop is regulated by TCR signaling strength and IL12. **A.** Left, CD69 expression on TCR_TAG_ stimulated *in vitro* with TAG altered peptide ligands (APL: low avidity TAG.D4 (blue), intermediate avidity TAG.F6 (green) and high avidity TAG.N4 (red) at concentrations ranging from 10^-3^-10^4^ nM. Summary plots show %CD69+, n=3 per group. Right, TCF1 expression on CD69+ TCR_TAG_ (gated as shown on left). Log_2_ FC of TCF1 MFI in comparison to no stimulation (0 nM) control. **B.** Naive TCR_OTI_ were stimulated for 12h with OVA APL: low avidity OVA.Q4 (blue), intermediate avidity OVA.A2 (green), and high avidity OVA.N4 (red). Left, CD69 expression with summary plots showing %69+ gated on CD8^+^ Thy1.1^+^ TCR_OTI_. Right, TCF1 expression on CD8^+^ Thy1.1^+^ CD69+ cells, summary plots show log_2_ FC of TCF1 MFI in comparison to no stimulation. \*\*\*\**P*<0.0001 determined by one-way ANOVA with post-hoc Tukey test. n=3 per group, representative of 2 independent experiments. **C, D**. CFSE-labeled naive TCR_OTI_ were stimulated with IL12 alone (black), high avidity OVA.N4 alone (light red) or in combination with IL12 (dark red), low avidity OVA.Q4 alone (light blue) or in combination with IL12 (dark blue). CFSE, and TCF1 expression are shown at 12h (C) or 72h (D) of stimulation. \**P*<0.05, \*\**P*=0.01, \*\*\**P*<0.001, \*\*\*\**P*<0.0001 determined by one-way ANOVA with post-hoc Tukey’s test shown for selected comparisons. n=3 per group, representative of 2 independent experiments.

Next, we wanted to determine the impact of inflammation on the pre-division TCF1 drop. Innate immune cells in *Listeria*-infected mice produce IL12^27^, which drives TCF1 downregulation as T cells proliferate^19^. High-avidity OVA.N4 peptide stimulation for 12h led to a pre-division TCF1 drop, which was not augmented by IL12 (**Fig. 2C**). However, IL12 treatment during low-avidity OVA.Q4 peptide-induced priming markedly potentiated the pre-division TCF1 drop (**Fig. 2C**). Interestingly, high-avidity OVA.N4 stimulation alone failed to induce the secondary proliferation-associated TCF1 drop (**Fig. 2D right**), while the addition of IL12 to both low- and high-avidity stimulation induced a robust proliferation-associated TCF1 drop (**Fig. 2D**). Interestingly, though T cells in liver tumors underwent the pre-division TCF1 drop followed by rebound, they did not exhibit the proliferation-associated secondary drop (**Extended Data** Fig. 1D). We previously showed that T cells in liver tumors are not exposed to IL12 or other pathogen-associated inflammatory cytokines, evidenced by their failure to activate inflammation-associated TF^6^. Thus, the pre-division TCF1 drop is regulated by both TCR signaling strength and inflammation, whereas the secondary TCF1 drop is primarily regulated by inflammation.

### Initial priming conditions determine effector versus memory differentiation

Though TCR signaling strength and inflammation are known to regulate T cell proliferation and differentiation^28,29^, we wanted to disentangle their role during the pre-division priming phase as opposed to during proliferation. Therefore, we asked whether initial pre-division priming conditions alone influence later CD8 T cell differentiation. For these studies, we utilized 1 nM OVA.N4 (N4) or OVA.Q4 (Q4) APL, which uniformly activate TCR_OTI_, as measured by CD69 upregulation (Fig. 2B left), but have a markedly differential ability to induce the pre-division TCF1 drop (Fig. 2B right). We stimulated congenically-distinct CFSE-labeled naive TCR_OTI_ *in vitro* with Q4 alone (minimal pre-division TCF1 drop) or Q4 in combination with IL12 (large pre-division TCF1 drop) for 12h, co-transferred the differentially-primed populations into B6 mice infected with empty LM (LM_Ø_), and harvested at early (2.5 days (d)) and late effector (5d) time points (**Fig. 3A**). By co-transferring differentially-primed T cells into the same host, where they experienced the same secondary inflammation, required for effector differentiation^30^, any differences observed can be solely attributed to the priming conditions. Low-avidity antigen during priming, with or without IL12, was sufficient to drive sustained proliferation (**Fig 3B left**). However, IL12 during priming led to preferential localization of differentiating T cells to tissue sites (liver) as compared to secondary lymphoid organs (spleen) (**Fig. 3B right**). Notably, IL12-primed TCR_OTI_ underwent a larger secondary TCF1 drop and had a larger population of cells expressing granzyme B (GZMB) and perforin (PRF1) (**Fig. 3C**). TCF1 and GZMB expression were inversely correlated (**Fig. 3C** right panels), as previously observed^31^. At the late effector time point (5d), when cells diverge between memory precursor cells (MPEC) and short-lived effector cells (SLEC)^30,32^, Q4 only-primed TCR_OTI_ maintained a larger stem/memory-like population (expressing TCF1 and CD62L^32^) than Q4+IL12 (**Fig. 3D**). We next tested the impact of TCR signal strength during priming by stimulating naive TCR_OTI_ with Q4 and N4 APL for 12h *in vitro* before transferring into LM_Ø_-infected B6 (**Extended Data** Fig. 2A). Stimulation with higher avidity peptide during priming led to greater effector differentiation (lower TCF1, higher GZMB) at early time points (**Extended Data** Fig. 2B) while lower avidity peptide stimulation promoted formation of more memory precursor-like (higher TCF1, lower GZMB, higher CD62L) (**Extended Data** Fig. 2C). Thus, TCR signaling strength and inflammation signaling during just the initial priming phase, which tunes the pre-division TCF1 drop, have a sustained impact on the secondary TCF1 drop and on effector versus memory differentiation.

**Figure 3.**
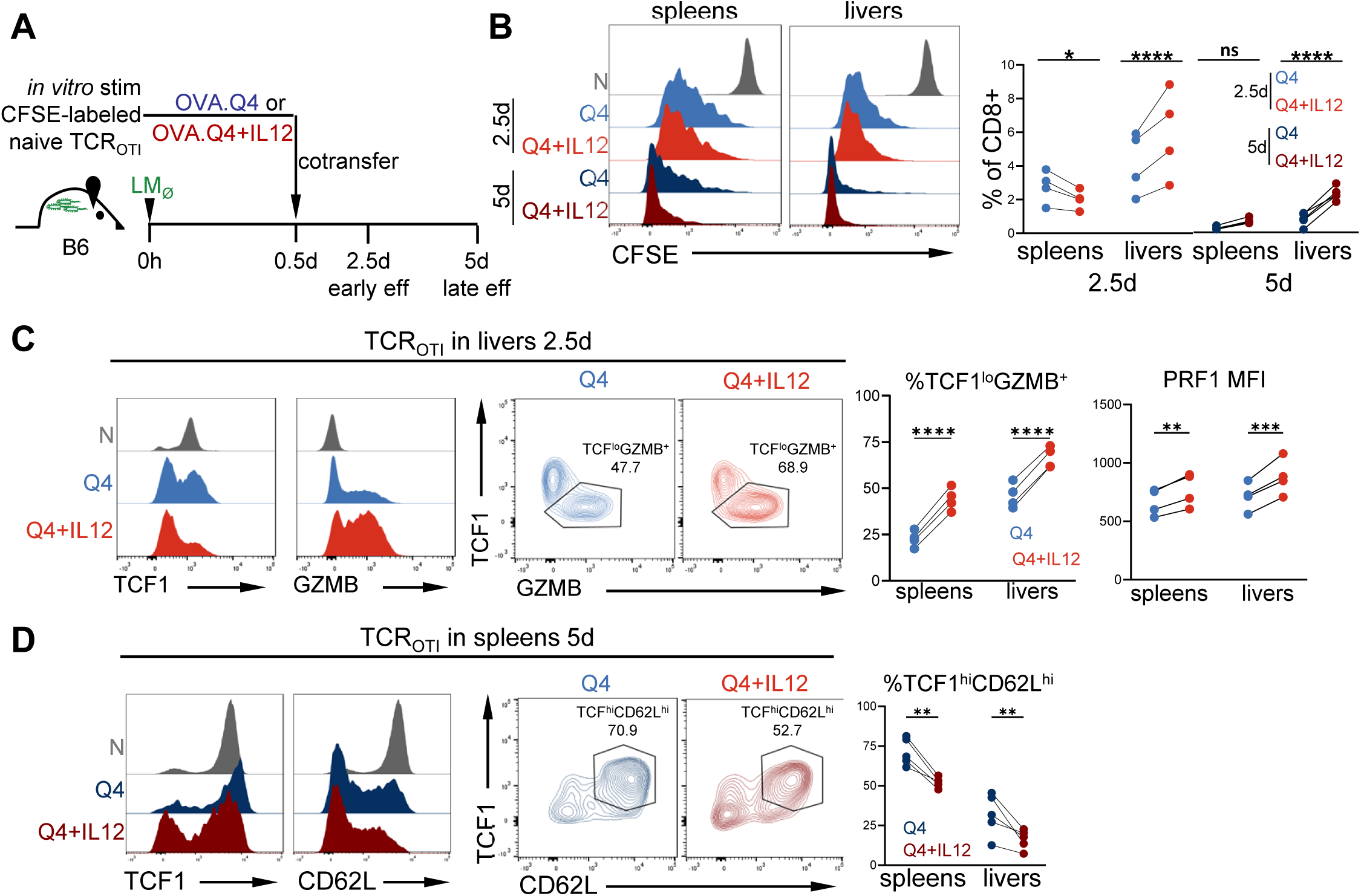
Initial priming conditions determine effector versus memory differentiation. **A**. Experimental scheme: CFSE-labeled naive TCR_OTI_ (Thy1.1 or Thy1.12) were stimulated *in vitro* with low avidity OVA.Q4 alone (blue) or OVA.Q4+IL-12 (red). After 12h, differentially primed T cells were co-adoptively transferred into LM_Ø_-infected B6 (Thy1.2). Lymphocytes were re-isolated from spleens and livers of infected mice and analyzed at indicated time points together with naive control (N;grey). **B.** Left, CFSE expression and right, % TCR_OTI_ of live CD8 T cells concatenated from 4 biologic replicates. \**P*=0.03, \*\*\*\**P*<0.0001 determined by two-way ANOVA with post-hoc Sidak multiple comparison test. n=4 per group, representative of two independent experiments. **C.** TCF1 and GZMB expression in the liver 2.5 days (d) following activation. Data was concatenated from 4 biologic replicates. TCF1^lo^GZMB^+^ gate set based on N. \*\**P*=0.0024, \*\*\**P*=0.0002, \*\*\*\**P*<0.0001 determined by two-way ANOVA with post-hoc Sidak multiple comparison test with n=4 per group. **D.** TCF1 and CD62L expression in the spleen 5d following activation. \*\**P*<0.01 determined by two-way ANOVA with post-hoc Sidak multiple comparison test. n=4-5 per group, representative of 2 independent experiments. Data was concatenated from 4 biologic replicates.

### Priming-encoded fates are not overridden by antigen re-encounter

T cells receive TCR and inflammatory signals not only during initial priming, but also during subsequent antigen encounters until pathogen clearance. We therefore investigated if priming encoded fate decisions are overridden by additional antigen encounters during the proliferation phase. We stimulated naive TCR_OTI_ with low (Q4) or high (N4) avidity peptide for 12h and transferred into B6 mice infected with LM encoding high-avidity N4 OVA peptide (LM_OVA_) (**Fig. 4A**). Remarkably, Q4 low-avidity primed T cells, despite being exposed to high-avidity antigen *in vivo*, generated a larger central memory population (TCF1^hi^CD62L^hi^) (30d time point) (**Fig. 4B**). In contrast, high avidity N4 priming generated a population with sustained enhanced effector function at 30d, as demonstrated by higher *in vitro* killing of OVA-pulsed target cells (**Fig. 4C left**). We re-challenged these mice again with a 2° LM_OVA_ infection to determine whether the initial priming impact would persist or be overridden. A greater proportion of N4-primed TCR_OTI_ were GZMB+ and had higher PRF1 as compared to Q4-primed TCR_OTI_ (32d after initial priming) (**Fig. 4C right)**. 5d after 2° infection, Q4 primed T cells had a larger TCF1^hi^CD62L^hi^ and CD127^hi^ KLRG1^lo^ population, reflecting a larger memory precursor population (**Fig. 4D**). We followed peripheral TCR_OTI_ numbers in 2° LM_OVA_-infected mice and found that while both groups underwent similar contraction over time (**Extended Data** Fig. 2D), the Q4 low avidity-primed group maintained a greater central memory TCF1^hi^CD62L^hi^ population, even 80d following 2° infection (**Fig. 4E**). The magnitude of the pre-division TCF1 drop, regulated by priming conditions, correlated with both the secondary proliferation-associated TCF1 drop and with effector versus memory differentiation. This raised the question of whether and how, the pre-division TCF1 drop determines long-term fate decisions.

**Figure 4.**
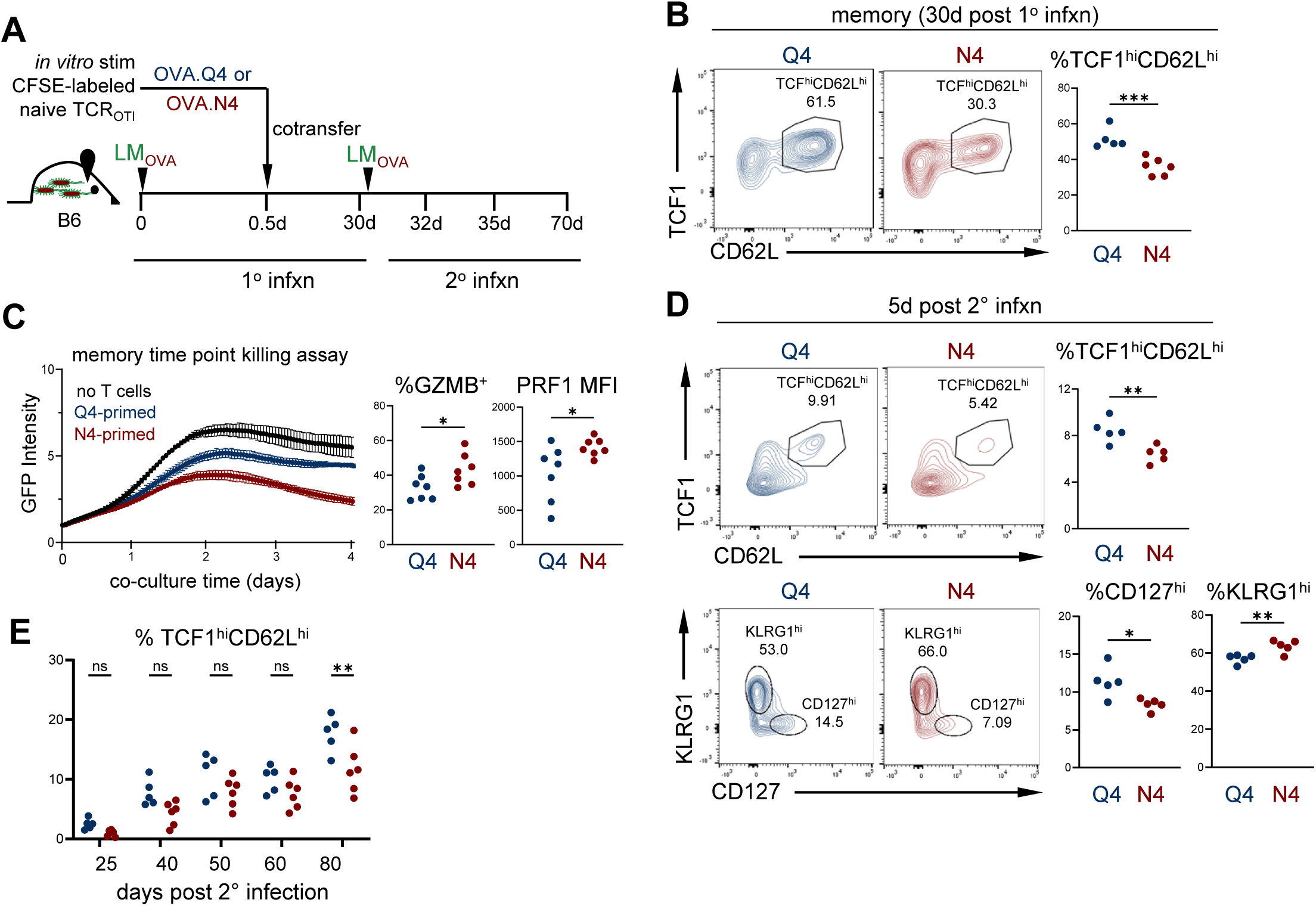
Priming-encoded fates are not overridden by antigen re-encounter. **A.** Experimental scheme: CFSE-labeled naive TCR_OTI_ (Thy1.12) were stimulated *in vitro* with low avidity OVA.Q4 (Q4;blue) or high avidity OVA.N4 (N4;red) and 12h later, adoptively transferred into LM_OVA_-infected B6 (Thy1.2). Mice were bled 30d following transfer. At a memory timepoint (30+d following transfer), mice were re-challenged with LM_OVA_. 2 and 5d later, lymphocytes were re-isolated from spleens and livers and analyzed. **B.** TCF1 and CD62L expression from TCR_OTI_ in the blood 30d post-transfer. P=0.0009 determined by unpaired t test. **C.** Left, at 30d, memory TCR_OTI_ were sorted from the spleen and cultured *in vitro* with EGFP+ tumor cell line (B16) pulsed with OVA.N4 peptide. Target cell killing was assessed by measuring EGFP intensity over 96h hours, normalized to the starting EGFP intensity Right, GZMB and PRF1 expression in the spleen 2d following rechallenge with LM_OVA_. \**P*<0.05 determined by unpaired t test. **D.** Top, CD62L by TCF1 expression and bottom, CD127 by KLRG1 expression in spleens 5d post 2° infection with LM_OVA_. \**P*<0.0164, \*\**P*<0.01 determined by unpaired t test. **E.** TCF1 and CD62L expression on TCR_OTI_ in the blood at indicated time points following 2° infection with LM_OVA_. \*\**P*=0.002 determined by two-way ANOVA with post-hoc Sidak multiple comparison test. n=5-7 per group, representative of 2 independent experiments (B-D).

### Pre-division TCF1 drop induces chromatin and transcriptional remodeling

As T cell differentiation is encoded through epigenetic and transcriptional changes, we investigated the impact of the pre-division TCF1 drop on T cell molecular programming. TCF1 controls gene expression in a complex manner, regulating chromatin accessibility and genomic architecture^33-35^ and through intrinsic histone deacetylase activity^36^, resulting in both transcriptional activation and repression^37^, We carried out RNA-sequencing (RNA-SEQ) paired with <u>A</u>ssay for <u>T</u>ransposase-<u>A</u>ccessible <u>C</u>hromatin-sequencing (ATAC-SEQ)^38^ on TCR_TAG_ differentiating over the course of LM_TAG_ infection. We sorted naive (CD44^lo^) and activated TCR_TAG_ from LM_TAG_-infected mice at pre-division (12h), early-division (48h), late effector (5d), and memory time points (30+d) (**Fig. 5A and** **Extended Data** Fig. 3A,B). For the early division (48h) time points, cells were sorted based on divisions 1-3 (P1-3), and P4-6 and CD25 (P4-6 CD25^lo^, P4-6 CD25^hi^), upregulated in cells undergoing effector differentiation ^18,39^. We analyzed these data together with our previously published data from TCR_TAG_ isolated from LM_TAG_-infected B6 mice at early (6-24h)^6^ and late time points (5d-30+d)^40^ to build a longitudinal map of chromatin and transcription landscape changes during acute infection. P4-6 CD25^hi^ and SLEC were TCF1^lo^, while P4-6 CD25^lo^, MPEC, and CM and EM were TCF1^hi^ (**Extended Data** Fig. 3C), and *Tcf7* mRNA expression closely followed TCF1 protein levels (**Extended Data** Fig. 3D). Principal component analysis (PCA) showed the greatest divergence in chromatin accessibility (**Extended Data** Fig. 4A) and transcription (**Extended Data** Fig. 4B) between naive and the E6h time points, with 20,182 differentially accessible chromatin peaks opening and 7,096 peaks closing (**Fig. 5B**). There was a second wave of chromatin remodeling as T cells transitioned from the pre-division to early division (P1-3) time points, and a third wave as T cells transitioned from effector/proliferative states to the memory time points (**Fig. 5B**).

**Figure 5.**
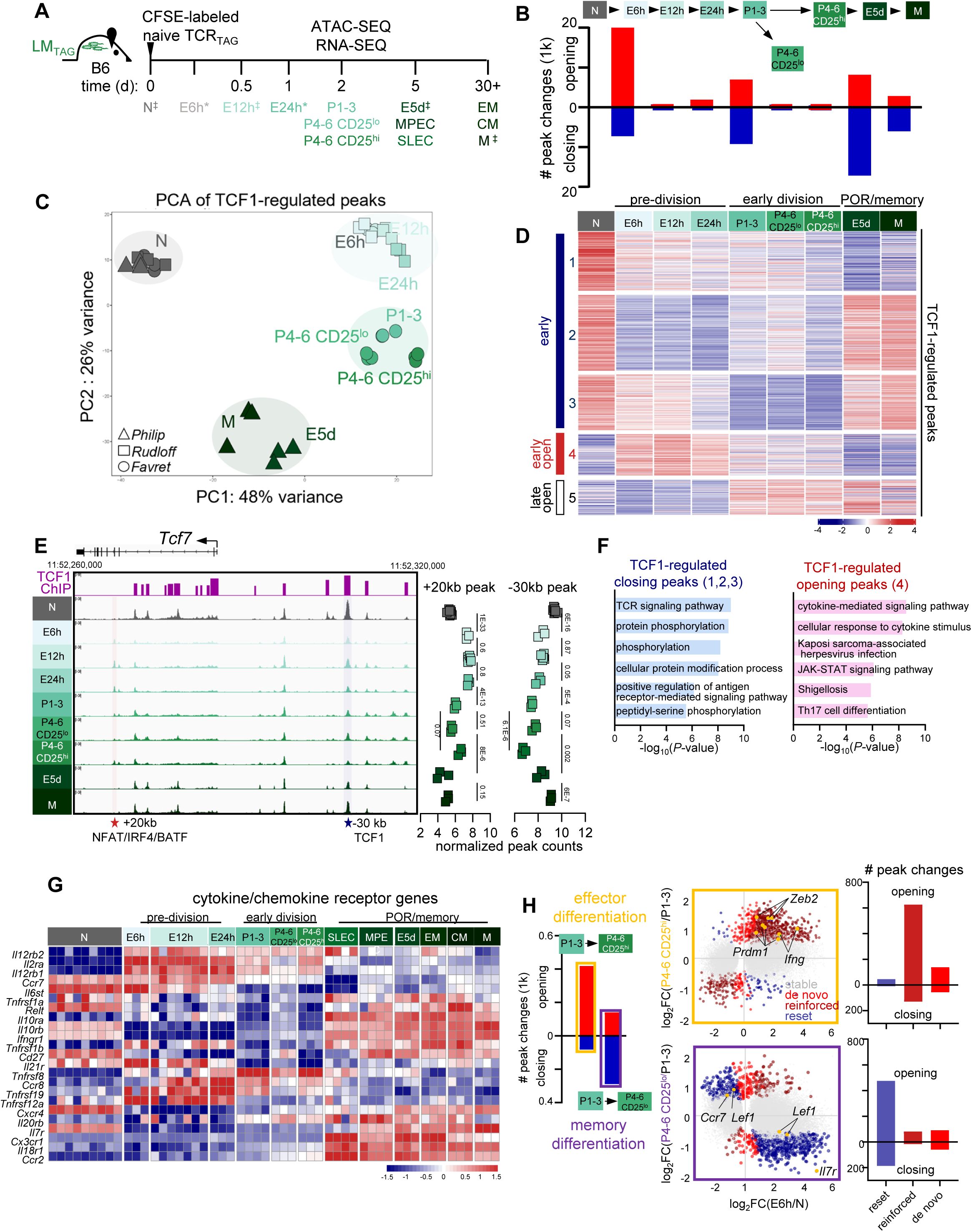
Pre-division TCF1 drop induces chromatin and transcriptional remodeling. **A.** Experimental/sorting scheme: CFSE-labeled naive TCR_TAG_(Thy1.1) were adoptively transferred into LM_TAG_-infected B6 mice; spleens were harvested at indicated time points and sorted as shown in **Extended Data** Fig. 3B. Time points labeled with the “*” (naive, 6h, 12h, 24h and 5d, 30+d) indicate that data was obtained from previously published studies (GSE209712)(GSE89308)^12,38^. Time points labeled with “^‡”^ indicate samples that were sorted in the current study and also represented in published data sets. **B.** Number of differentially accessible chromatin peaks (DAC) changing during each transition (opening peaks, red; closing peaks, blue; false discovery rate (*FDR*)<0.05; absolute log_2_foldchange > 1). **C.** PCA comparing chromatin accessibility of TCF1 ChIP+ motif+ peaks in naive and differentiated CD8 T cells. **D.** Heatmap of TCF1 ChIP+ motif+ peaks clustered based on supervised k-Means (k=5) by the direction of peak change between N to the pre-division time points and z-score normalized across rows. **E**. Chromatin accessibility profile across the *Tcf7* locus. Purple bars at the top indicate TCF1 ChIP binding sites^34^ (GSE164670). +20kb peak (red star) and quantified (right) indicates a negative regulatory site shown to be a binding site for transcription factors NFAT, IRF4 and BATF. -30kb peak (blue star) and quantified (right) indicates positive regulatory site bound by TCF1. *FDR* values as determined by DEseq2 shown for sequential time points. **F.** EnrichKG pathway analysis of TCF1-regulated closing peaks (left) and opening peaks (right). **G.** Heatmap showing cytokine/chemokine receptor genes (KEGG map04060) containing TCF1-regulated peaks that were differentially-expressed by LRT analysis (*P*<0.0001), z-score normalization across rows. **H.** Left, number of differentially accessible chromatin peaks (DAC) changing during P1-3->P4-6 CD25^hi^ (yellow box) and P1-3->P4-6 CD25^lo^ (blue box) transitions. Right upper, DAC during naive (N) → early (E6h) transition (log_2_FC E6h/N) versus P1-3 → P4-6 CD25^hi^ (E5 d) transition (log_2_FC P1-3/ P4-6CD25^hi^). Lower, DAC during N → early (E6h) transition (log_2_FC E6h/N) versus P1-3 → P4-6 CD25^lo^ transition (log_2_FC P1-3/ P4-6 CD25^lo^). Each point represents an individual peak colored as in legend (grey=stable (unchanging in P1-3->P4-6 transition), red=de novo (changing only during P1-3->P4-6 transition), dark red=reinforced (changes during N->E6h transition and change in the same direction during P1-3->P4-6 transition), blue=reset (changes during N->E6h transition changes in the opposite direction during P1-3->P4-6 transition). FC determined by DESeq2 analysis, *FDR*<0.05. Right, bar plots show the number of de novo peaks in the reset, reinforced, and de novo categories, *FDR* < 0.05 for each comparison.

We used published TCF1 ChIP-SEQ seq data^37^ and identified 2,419 peaks with TCF1 binding by ChIP and containing the TCF1 DNA binding motif (JASPAR MA0769.3^41^), indicating high-confidence sites of TCF1 direct binding and regulation^37^. Interestingly, PCA of TCF1-regulated chromatin peaks showed that T cells clustered by their time point/differentiation stage rather than TCF1 expression (**Fig. 5C**). Therefore, we compared how TCF1-regulated peaks changed over time, grouping peaks into clusters by their patterns of accessibility (**Fig. 5D**). Strikingly, most TCF1-regulated peaks (clusters 1-3: 1726/2419, ∼70%) decreased in accessibility early after activation (E6-24h) (**Fig. 5D**), coinciding with the pre-division drop in TCF1. Intriguingly, these peaks remained closed at the later post-division time points (48h) despite the rebound in TCF1 protein in the P1-3 and P4-6 CD25^lo^ groups (**Fig. 5D**, **Extended Data** Fig. 3C).

Twenty differentially accessible peaks were found within the *Tcf7* locus itself (**Fig. 5E**). Consistent with our finding that TCR signal strength impacted the magnitude of the TCF1 drop (Fig. 2), we identified a known repressive peak at +20 kb (**Fig. 5E**) that was previously shown to bind TCR signaling-induced TF (NFAT, IRF4 and BATF)^42^. Indeed this peak increased in accessibility during pre-division time points (**Fig. 5E right**), suggesting that TCF1 signaling-activated TF repress *Tcf7* expression. Several peaks within the *Tcf7* locus and upstream regulatory regions^43^ contained TCF1 binding sites (**Fig. 5E**), including a peak at -30 kb with high TCF1 binding and a TCF1 binding motif, which decreased in accessibility at the pre-division time points, consistent with TCF1’s known autoregulation^44^. Taken together, these findings suggest a temporally-ordered TF network in which TCR-induced TF bind repressive peaks in the *Tcf7* locus, downregulating *Tcf7*/TCF1 expression and impeding TCF1 autoregulatory expression. To understand the biologic consequences of the widespread pre-division TCF1-mediated chromatin closing, we ran EnrichrKG pathway analysis^45^ on genes containing closing peaks (clusters 1, 2, 3). In these early closing peaks, we found enrichment for TCR signaling pathways (**Fig. 5F left**). Accordingly, genes encoding TCR signaling proteins were downregulated at the early pre-division time points (*Lck*, *Cd28*), and among the few upregulated genes were several negative regulators of TCR signaling (*Ctla4*, *Cblb*, *Ppp2cb*) (**Extended Data** Fig. 4D). Thus, physiologic TCR negative feedback pathways are engaged early after activation, mediated in part by the pre-division TCF1 drop.

Our analysis of the smaller group of TCF1-regulated peaks which opened pre-division and remained open at 48h (cluster 4: 374/2419, 15%) revealed an interesting finding: these peaks were enriched for pathways associated with cytokine-mediated signaling (**Fig. 5F right**). Indeed, mRNA for genes encoding receptors for inflammatory cytokines receptor (*Il12rb1*, *Il12rb2*, and *Il2ra*) was upregulated within 6h of activation (**Fig. 5G**), consistent with previous work showing that IL12RB is upregulated in *Tcf7*^-/-^ T cells^19^, and suggesting that the pre-division TCF1 drop may poise T cells to receive additional IL12 stimulation during the proliferation phase. Among peaks that opened early in response to the pre-division TCF1 drop (cluster 4) were peaks in the effector gene *Gzmb*. Peaks with ChIP-demonstrated TCF1-binding in the *Gzmb* locus increased in accessibility with the pre-division TCF1 drop, accompanied by increased Gzmb mRNA (**Extended Data** Fig. 4C), making these likely TCF1-repressed sites. Thus, the pre-division TCF1 drop promotes chromatin remodeling and expression or poising of genes encoding effector molecules and inflammatory cytokine receptors.

Peaks that did not change during the pre-division phase (cluster 5: 319/2,419, 13%) gained accessibility later as T cells proliferated (**Fig. 5D**), including peaks in genes *Gzma*, *Fasl*, and *Runx3*. To understand how the pre-division chromatin accessibility evolved over time, we assessed chromatin peak changes during the early proliferation phase. Interestingly, as P1-3 divided and became P4-6 CD25^hi^ TCF1^lo^ early effectors, many chromatin peaks opened (**Fig. 5H left**). In contrast, the transition to memory-precursor-like P4-6 CD25^lo^ TCF1^hi^ state entailed mainly peak closing (**Fig. 5H left**).To understand this difference, we analyzed individual chromatin peak accessibility during the pre-division transition (N->E6h) as compared to the early proliferation transition (P1-3->P4-6 CD25^hi^ or P1-3->P4-6 CD25^lo^). Many peaks did not change during the later transition (stable; grey); however, among the changing peaks we observed a striking difference (**Fig. 5H**). As T cells underwent effector differentiation (CD25^hi^), pre-division peak changes (N->E6h) were reinforced, that is peaks that gained accessibility pre-division had additional gain in accessibility with proliferation, while peaks that lost accessibility pre-division closed further (755/1,005 changing peaks, 75%) (**Fig. 5H upper middle and right**). These reinforced peaks were in known effector-associated genes, including *Zeb2*, *Ifng*, and *Prdm1*^46,47^. On the other hand, as T cells proliferated and underwent memory differentiation, DAC were reset (e.g. peaks opening pre-division closed with proliferation and vice versa; 662/911 changing peaks, 73%), including peaks in naive/memory-associated genes such as *Lef1*, *Il7r, Ccr7*^30,34,37^ (**Fig. 5H lower middle and right**). There were fewer *de novo* peak accessibility changes during the transition from P1-3 to P4-6 during either effector (195/1005, 19%) or memory (151/911, 17%) differentiation (**Fig. 5H right**). Taken together, our epigenetic and transcriptional analyses demonstrate that the pre-division TCF1 drop remodels the T cell epigenome to poise effector gene expression and increase sensitivity to inflammatory cytokines, providing a means for early priming conditions to impact later differentiation.

### Pre-division TCF1 drop enables inflammation-driven effector differentiation

Since we observed that TCF1-repressed cytokine receptors such as IL12R were upregulated during the pre-division TCF1 drop, we hypothesized that TCF1 tuning during priming controls inflammatory cytokine-driven effector differentiation. To test this, we transiently transfected T cells with siRNA targeting *Tcf7* to downregulate TCF1 during the pre-division phase independently of priming conditions. CFSE-labeled TCR_OTI_ were transfected with anti-*Tcf7* siRNA (TCF1kd) or non-targeting control siRNA (ntc), rested, and primed with low-avidity Q4 peptide (**Fig. 6A**). TCF1kd cells had lower TCF1 expression prior to stimulation (0h), which dropped further with Q4 stimulation, while the ntc remained TCF1+ and dropped minimally with stimulation (**Fig. 6B, C**). Both the TCF1kd and ntc underwent the TCF1 rebound on cell cycle entry (**Fig. 6B, C**). Transient TCF1 knockdown did not impact early activation, as shown by similar CD69 upregulation and CD62L downregulation (**Extended Data** Fig. 5A) and proliferation/cell numbers compared to ntc (**Extended Data** Fig. 5B). We restimulated *in vitro* with Q4, IL12, or Q12+IL12 and found that transient pre-division TCF1kd impacted the secondary TCF1 drop in response to IL12 or Q4+IL12, but not to Q4 alone (**Fig. 6D**), consistent with our finding that early epigenetic remodeling led to an increase in IL12R expression (**Fig. 5G**).

**Figure 6.**
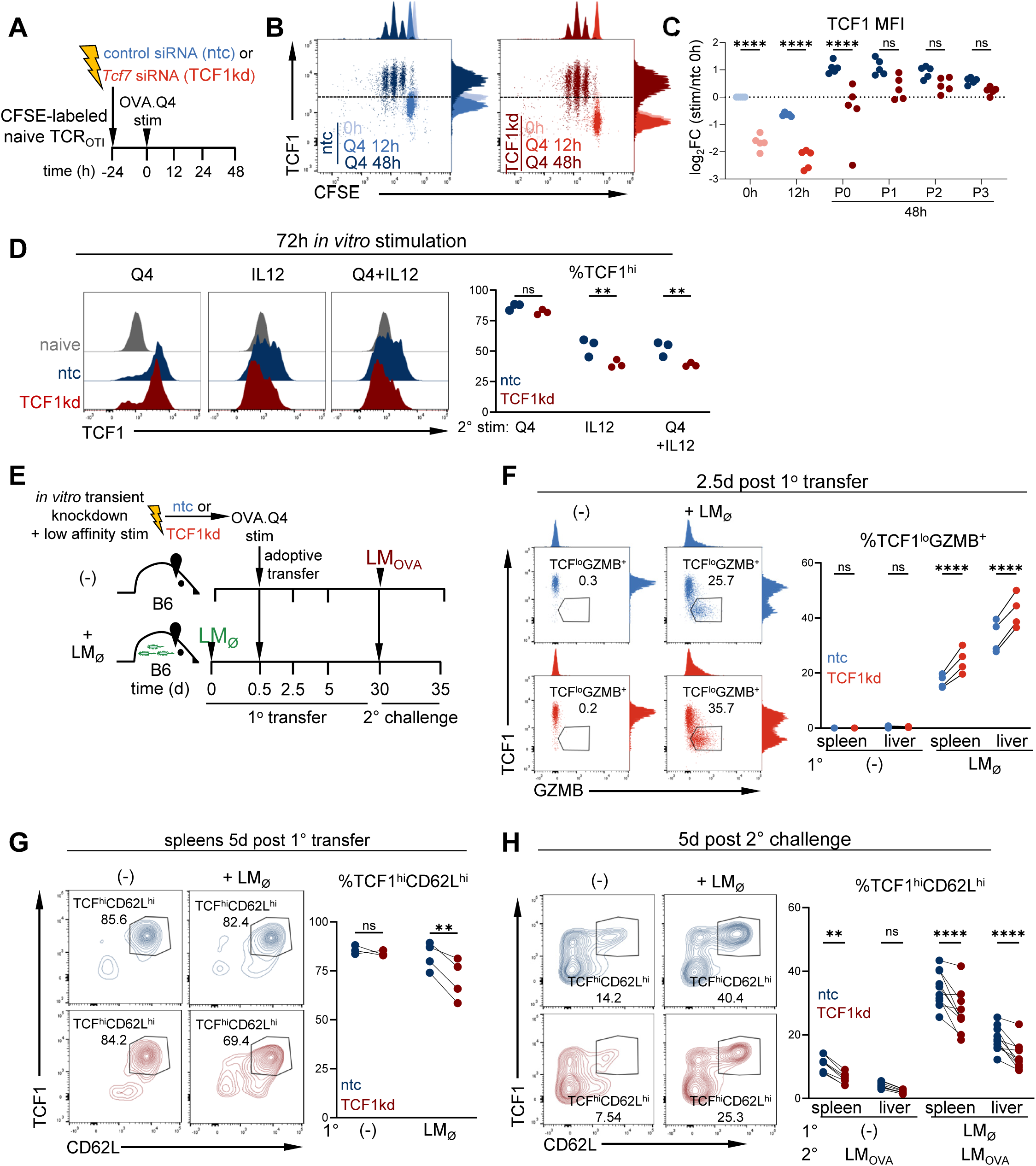
Pre-division TCF1 drop enables inflammation-driven effector differentiation. **A.**, Experimental scheme: CFSE-labeled naive TCR_OTI_ (Thy1.12) were electroporated with nontargeting control (ntc:blue) or *Tcf7* targeting (TCF1kd:red) siRNA, rested, stimulated *in vitro* with low avidity Q4, and analyzed at indicated time points. **B.** CFSE by TCF1 expression at indicated timepoints. **C.** Summary plot of TCF1 expression in ntc vs TCF1kd at indicated time points/division. Log_2_ FC of TCF1 MFI of TCR_OTI_ in comparison to 0h ntc. \*\*\*\**P*<0.0001 determined by two-way ANOVA with post-hoc Sidak multiple comparison test. n=3-5 per group, representative of two independent experiments. **D.** Naive TCR_OTI_ were transfected with siRNA and stimulated with Q4 *in vitro*; at 48h, ntc (blue) and TCF1kd (red) cells were re-stimulated with Q4 (low avidity), IL12, or a combination of Q4+IL12. At 72h (24h following restimulation), cells were assessed for TCF1 expression. TCF1^hi^ was gated based on no stim ntc control. \**P*=0.05, \*\**P*=0.0017 determined by two-way ANOVA with post-hoc Sidak multiple comparison test. n=3 per group, representative of two independent experiments. **E.** Experimental scheme: CFSE-labeled naive TCR_OTI_ were electroporated with nontargeting control (ntc:blue) or *Tcf7* targeting (TCF1kd:red) siRNA, rested, then stimulated *in vitro* with low avidity OVA.Q4. At 12 hours, T cells were adoptively co-transferred into uninfected or LM_Ø_-infected B6. 2.5 and 5d following transfer, lymphocytes were re-isolated from spleens and livers and analyzed by flow cytometry. At a memory timepoint (21+d following transfer), mice were re-challenged with LM_OVA_. 5d later, lymphocytes were re-isolated from spleens and livers and analyzed by flow cytometry. **F.** GZMB by TCF1 expression 2.5 days following activation/transfer. \*\*\*\**P*<0.0001, determined by two-way ANOVA with post-hoc Sidak multiple comparison test. **G.** TCF1 by CD62L expression 5d post-transfer. \*\**P*=0.004, determined by two-way ANOVA with post-hoc Sidak multiple comparison test. n=4 per group, representative of two independent experiments. **H.** TCF1 by CD62L expression 5d after 2° challenge. \*\**P*=0.002, \*\*\**P*<0.0001 determined by two-way ANOVA with post-hoc Sidak multiple comparison test. n=8-10 per group, representative of two independent experiments.

To test the impact of early TCF1 tuning on *in vivo* inflammation-driven differentiation and long-term fate commitment, we co-transferred Q4-primed ntc and TCF1kd TCR_OTI_ into uninfected or LM_Ø_-infected B6 and assessed differentiation at early (2.5d) and late effector/memory precursor (5d) time points (**Fig. 6E**). At the early effector time point TCF1kd cells gained greater GZMB (**Fig. 6F**) and PRF1 (**Extended Data** Fig. 5C) expression and showed increased migration to the liver in LM_Ø_-infected recipients, while neither TCF1kd nor ntc cells underwent effector differentiation in B6 mice (**Fig. 6F**, **Extended Data** Fig. 5C,D). At the late effector time points, more TCF1kd than ntc downregulated TCF1 and CD62L in LM_Ø_-infected B6, but not in uninfected recipients (**Fig. 6G**). Thus, TCF1 tuning during priming impacts the degree of effector differentiation in response to inflammatory signaling, but not in the absence of inflammation. We next wanted to determine whether the impact of transient priming-induced TCF1 tuning would persist after 2° challenge with both antigen and inflammation (LM_OVA_). Prior to re-challenge with LM_OVA_, both ntc and TCF1kd had similar numbers of TCR_OTI_ in the peripheral blood (**Extended Data** Fig. 5E), and following re-challenge both TCF1kd and ntc expanded robustly (**Extended Data** Fig. 5F). Strikingly, TCF1kd again underwent greater effector differentiation (decreased CD62L) and less memory precursor differentiation (decreased TCF1, decreased EOMES) compared to ntc, even after 2° LM challenge (**Fig. 6H**, **Extended Data** Fig. 5G). Thus, the pre-division TCF1 drop “imprints” T cell fates, skewing the proliferating T cell population towards greater effector differentiation in response to inflammatory cytokines (**Extended Data** Fig. 5H).

## DISCUSSION

We show here for the first time that in CD8 T cells encountering antigen during infection and cancer, TCF1 is dynamically regulated during priming prior to cell division. *Tcf7* mRNA is rapidly downregulated within hours of antigen encounter in both acute infection and in solid tumors. As TCF1 protein is continuously translated and degraded in T cells^22^, *Tcf7* transcriptional downregulation results in rapid loss of TCF1 protein. The magnitude of the pre-division drop is regulated by TCR signal strength and inflammatory cytokines. TCF1 expression rebounds with cell cycle entry and declines again as CD8 T cells proliferate and undergo effector differentiation (secondary drop). Remarkably, transient siRNA-mediated TCF1 downregulation during the pre-division priming phase was sufficient to bias long-term differentiation toward the effector fate. This pre-division TCF1 drop and rebound is a previously unrecognized step in TCF1 regulation, begging the question of how and why it occurs.

While effector differentiation was thought to occur as T cells proliferate, we demonstrate that within 6h of priming initiation, activated CD8 T cells undergo large-scale chromatin remodeling, including in TCF1-regulated genes, which dictates long-term fate commitment. Upon pre-division priming and TCF1 downregulation, many TCF1-regulated inhibitory and activating peaks underwent remodeling, including peaks within the *Tcf7* locus. Chromatin remodeling in TCF1-inhibited genes caused the activation and expression of genes encoding inflammatory cytokine receptors, such as *Il12rb2* and effector molecules such as *Gzmb*. While the greatest number of chromatin peak changes occurred upon priming pre-division, poising activated CD8 T cells for effector function, as T cells began proliferating and committed to the effector fate (CD25^hi^) upregulation, additional peak changes reinforced the pre-division accessibility changes (in genes such as *Zeb2*, *Ifng*, and *Prdm1*). In contrast, proliferating CD25^lo^ T cells (memory precursors) underwent chromatin remodeling that reset the pre-division accessibility changes to a more naive-like state (including in genes such as *Lef1*, *Il7r*, and *Ccr7*). This chromatin accessibility reset we observed is in line with previous work showing that memory precursor cells undergo erasure of activation-induced DNA methylation in naive/memory genes^48^.

Together, our findings suggest that the pre-division TCF1 drop functions as an imprinting mechanism, linking priming conditions to long-term epigenetic and transcriptional programs that ensure CD8 T cell fates remain tuned to the surrounding environment. During acute infection, initial priming results in a population of activated T cells that have undergone varying degrees of TCF1 downregulation, determined by their antigen avidity and the cytokine milieu. Thus, after priming, activated T cells will be heterogeneous with regards to their imprinted long-term effector versus memory fate. As long as pathogen and associated inflammatory cytokines are present, effector-skewed T cells will proliferate, undergo the secondary TCF1 drop, and commit to the effector fate. On the other hand, if pathogen is rapidly cleared and there is little inflammatory cytokine, T cells will stop proliferating, contract, and/or undergo memory differentiation. This mechanism of priming-induced pre-division fate specification allows the responding CD8 T cell population to scale the number of effector and memory T cell precursors early during infection, while maintaining flexibility to generate more or fewer effectors as required over the course of infection.

While the pre-division TCF1 drop poises T cells for the effector fate, the role of the TCF1 rebound is less clear. One possibility is that the TCF1 rebound serves to maintain memory fate flexibility. TCF1 is required to maintain a central memory population capable of responding to secondary infectious challenges^15,49-51^, and the TCF1 rebound could increase the pool of memory precursors. This is in line with a recent study showing that flexibility in TCF1 expression confers scalability to the memory population size^52^. Another intriguing hypothesis is that the TCF1 rebound acts as a second checkpoint for the acquisition of effector function, thereby preventing autoimmunity. T cells encountering self-antigen in secondary lymphoid tissue (for example as a result of molecular mimicry with pathogen) might undergo the pre-division TCF1 drop. However, following T cell migration to the periphery, T cells would not undergo the secondary TCF1 drop and effector differentiation unless inflammatory cytokines were also present in the target tissue. Thus, the stepwise regulation of TCF1 may serve as a failsafe to prevent improper effector differentiation by requiring inflammation to be present at multiple stages of differentiation. However, the existence of multiple effector checkpoints may hamper anti-tumor responses when tumor-specific T cells activated in secondary lymphoid organs do not receive pathogen-associated cytokine signaling when they subsequently encounter antigens in tumors and thus fail to differentiate into effectors^6^.

TCF1 is a pivotal regulator of T cell development, differentiation, and function across the T cell life cycle, from the thymus to secondary lymphoid organs to the periphery. Now we reveal a regulatory role for TCF1 in the earliest hours following T cell priming, which impacts long-term T cell differentiation. Our study establishes the importance of the pre-division priming phase and TCF1 regulation for T cell fate determination and has potential therapeutic implications for T cell responses during infection, autoimmunity, and cancer therapy.

## METHODS

### Mice

TCR_TAG_ transgenic mice (B6.Cg-Tg(TcraY1,TcrbY1)416Tev/J) ^17^, TCR-OT1 (C57BL/6-Tg(TcraTcrb)1100Mjb/J) ^53^, Ly5.1 (C57BL/6J-Ptprcem6Lutzy/J) and C57BL/6J Thy1.1 mice were purchased from The Jackson Laboratory. TCR_TAG_;Thy1.1 double transgenic mice were generated by crossing Thy1.1 mice to TCR_TAG_ mice. TCR_OTI_;Ly5.1 double transgenic mice were generated by crossing the TCR-OTI mice with Ly5.1 mice. ASTxAlb-Cre^54^ double transgenic mice were generated by crossing AST (Albumin-floxStop-SV40 large T antigen (TAG))^55^ with Alb-Cre mice. ASTxAlb-Cre were used between 9 and 12 weeks of age, at which time all mice had established liver tumors. Both female and male mice were used for studies. T cell donor mice were between 6-10 weeks of age and sex-matched to recipient male and female C57BL/6 recipients. All mice were bred and housed in the animal facility at Vanderbilt University Medical Center (VUMC). All animal experiments were performed in compliance with VUMC Institutional Animal Care and Use Committee (IACUC) regulations.

### Adoptive T cell transfer of naive or primed cells in acute infection and tumor models

C57BL/6 mice were inoculated i.v. with 5-9 x10^6^ CFU of either *Listeria monocytogenes* (Lm) Δ*actA* Δ*inlB* strain ^56^, referred to as LM _Ø_, Lm expressing the TAG-I epitope (SAINNYAQKL, SV40 large T antigen 206–215), referred to as LM_TAG_ or the OVA epitope (SIINFEKL) (Aduro Biotech), referred to as LM_OVA_, 6-12h prior to T cell adoptive transfer for generation of effectors. Spleens from naive transgenic mice were mechanically disrupted with the back of 3 mL syringe and filtered through a 70 mm strainer into ammonium chloride potassium (ACK) buffer to lyse erythrocytes. For naive T cell transfers, cells were washed twice with cold serum-free RPMI 1640 media and 2.5x10^6^ TCR_TAG_;Thy1.1 CD8^+^ T cells were adoptively transferred into C57BL/6 (Thy1.2) mice inoculated with LM_TAG_. For priming, cells were resuspended in cold RPMI 1640 supplemented with 2 μM glutamine, 100 U/mL penicillin/streptomycin, and 10% FBS (cRPMI) + 2mercaptoethanol + OVA.N4 or OVA.Q4 peptide (1nM) or peptide + IL12 (20ng/ml) and incubated for 12 hours at 37°C. Cells were then washed with cold serum-free RPMI and either 1x10^5^ or 5x10^5^ activated TCR_OTI_;Thy1.12 or Ly5.12 CD8^+^ T cells were adoptively transferred into C57BL/6 (Thy1.2) mice inoculated with LM_OVA_ or LM_Ø_, respectively. For CFSE labeling studies, splenocytes were resuspended after first wash in 2.5 mL of plain, serum-free RPMI 1640, rapidly mixed with equal volumes of 2x CFSE [10mM] solution, incubated for 5 min at 37°C at a final CFSE [5 μM], quenched by mixing CFSE/cell solution with equal volume of pure FBS, washed twice with serum-free RPMI, and resuspended in serum-free RPMI for transfer.

### T Cell Isolation and Electroporation/Transfection

Spleens from naive transgenic mice were mechanically disrupted with the back of 3 mL syringe and filtered through a 70 mm strainer into 2%FBS/PBS buffer. Naïve CD8 T cells were magnetically isolated using EasySep^TM^ Mouse Naïve CD8+ T Cell Isolation Kit (STEMCELL Technologies). T cells were then counted, spun down, and resuspended in Opti-Mem. Cell suspension was mixed with either siGENOME nontargeting siRNA control Pools, Pool #1 (Horizon/Dharmacon) or mouse Tcf7 targeting siGENOME SMARTPool siRNA (Horizon/Dharmacon) and added to a 2mm sterile cuvette (Bulldog Bio). siRNA was reconstituted in nuclease-free water as directed (Dharmacon protocol). Cells were then electroporated using NEPA21 Electro-Kinetic Transfection System (Bulldog Bio). Immediately after electroporation, pre-warmed cRPMI containing 100ng/ml of IL-7 was added to the cuvette, and cuvette was then incubated at 37C for 15 minutes. Then, cells were gently mixed by pipetting and added to a flat-bottom 48-well plate. Additional warmed cRPMI was used to wash cuvette and added to well. Cells were rested for 8-10 hours following electroporation and then combined with splenocytes (no stim control) or stimulated with splenocytes and OVA.Q4 peptide. For *in vivo* transfers, cells were collected 12 hours following stimulation. Cells were then washed with cold serum-free RPMI and 3x10^5^ activated TCR_OTI_ were adoptively transferred into C57BL/6 (Thy1.2) mice inoculated with LM_Ø_, respectively. For *in vitro* analysis, cells were re-stimulated with OVA.Q4 peptide, IL12, or a combination of OVA.Q4 and IL12 and then analyzed 24 hours following re-stimulation

### Cell isolation for subsequent analyses

Spleens from experimental mice were mechanically disrupted with the back of 3 mL syringe and filtered through a 70 mm strainer into ACK buffer. Cells were washed once and resuspended in cold cRPMI. Liver tissue was mechanically disrupted using a 150 mm metal mesh and glass pestle in ice-cold 2% FBS/PBS and passed through a 70 mm strainer. Liver homogenate was centrifuged at 400g for 5 min at 4°C and supernatant discarded. Liver pellet was resuspended in 15 mL of 2% FBS/PBS buffer containing 500 U heparin, mixed with 10 mL of Percoll (GE) by inversion, and centrifuged at 500g for 10 min at 4°C. Supernatant was discarded and pellet was RBC lysed in ACK buffer and resuspended in cRPMI for downstream applications.

### Human PBMC isolation and culturing

PBMCs were isolated from healthy donor blood samples (Gulf Coast Regional Blood Center)using Sepmate-50 tubes (Stem Cell Technologies, catalog #1111587) and Ficoll-paque (Fisher, catalog #45-001-749). Cells were then lysed in ACK lysis buffer, resuspended in RPMI, and counted. For CFSE labeling studies, splenocytes were resuspended after first wash in 2.5 mL of plain, serum-free RPMI 1640, rapidly mixed with equal volumes of 2x CFSE [10mM] solution, incubated for 5 min at 37°C at a final CFSE [5 μM], quenched by mixing CFSE/cell solution with equal volume of pure FBS, washed twice with serum-free RPMI, and resuspended in cRPMI with IL-7 for no stimulation or cRPMI with aCD3, aCD28, and IL2 for stimulation and then incubated at 37C.

### Transcription factor staining

Transcription factor staining was performed with the Foxp3/Transcription Factor Staining Buffer Kit (Tonbo) per manufacturer’s instructions. Briefly, cells were stained with for surface markers, fixed, permeabilized, and stained for transcription factors or intracellular proteins.

### Flow cytometry and flow sorting

All flow analysis was performed on the Attune NXT Acoustic Focusing Cytometer (ThermoFisher Scientific). Data was analyzed using FlowJo v.10.10.0 (Tree Star Inc.). Cell sorting was performed using the BD FACS Aria III (BD Biosciences) at the VUMC Flow Cytometry Shared Resource Core with BD FACSDiva Software.

### Incucyte/In vitro Killing Assay

B16-EGFP were pulsed with OVA.N4 peptide overnight, cultured in a flat bottom 96-well plate in cRPMI. TCR_OTI_ sorted from spleens at memory timepoints and combined (n=9 per group), 2 technical replicates were cultured with B16 cells and then imaged every hour for 96 hours. Total fluorescence intensity normalized to first image/hour 0 intensity.

### RNA sequencing (RNA-SEQ)

ACK lysed single cell suspensions from livers and spleens were processed as described above using sterile technique and stained with antibodies against CD8, CD90.1, and CD69 and (4’,6-diamidino-2-phenylindole) DAPI for dead cell exclusion. 25,000 cells were sorted directly into Trizol LS and frozen. Total RNA was extracted from sorted cells using the Rneasy Micro kit (Qiagen) and amplified using the SMART-Seq v4 UltraLow Input RNA Kit (Clontech). The cDNA was quantified and analyzed on the BioAnalyzer. Libraries were prepared using 7.7-300 ng of cDNA and the NEB DNA Ultra II kit. Each library was quantitated post PCR and run on the Caliper GX to assess each library profile. A final quality control assay consisting of qPCR was completed for each sample. The libraries were sequenced using the NovaSeq 6000 with 150 bp paired end reads targeting 50M reads per sample. RTA (version 2.4.11; Illumina) was used for base calling and analysis was completed using MultiQC v1.7.5.

### ATAC sequencing (ATAC-SEQ)

Profiling of chromatin was performed by ATAC-seq as previously described ^38^. ACK lysed single cell suspensions from livers and spleens were processed as described above using sterile technique and stained with antibodies against CD8, CD90.1, and CD69 and (4’,6-diamidino-2-phenylindole) DAPI for dead cell exclusion. 25,000 cells were sorted into cold FCS, DMSO added to 10%, and cells frozen. Frozen T cells were then thawed and washed in cold PBS and lysed. The transposition reaction was incubated at 42°C for 45 min. The DNA was cleaned with the MinElute PCR Purification Kit (Qiagen), and material was amplified for five cycles. After evaluation by real-time PCR, 7–13 additional PCR cycles were done. The final product was cleaned by AMPure XP beads (Beckman Coulter) at a 1× ratio, and size selection was performed at a 0.5× ratio. Libraries were sequenced on a HiSeq 2500 or HiSeq 4000 in a 50-bp/50-bp paired-end run using the TruSeq SBS Kit v4, HiSeq Rapid SBS Kit v2, or HiSeq 3000/4000 SBS Kit (Illumina).

### Statistics and reproducibility

No statistical method was used to predetermine sample size, but sample sizes are similar to those reported in previous publications^6^. No data were excluded from the analyses. The experiments were not randomized. The investigators were not blinded to allocation during experiments and outcome assessment. Statistical analysis of next-generation sequencing data is described in detail above. All other experiments and statistical analysis, including two-tailed Student’s *t*-test, one-way ANOVA with post hoc Tukey test, two-way ANOVA with post hoc Sidak test, and one-sample Student’s *t*-test, were performed as described using Prism 9.0 (GraphPad). Data distribution was assumed to be normal, but this was not formally tested. Composite figures and schematics were generated using Powerpoint 365 (Microsoft).

### Bioinformatics methods

The quality of the sequenced reads was assessed with FastQC ^57^ and QoRTs ^58^ (for RNA-seq samples). Unless otherwise stated, plots involving high-throughput sequencing data were created using R v4.4.2 ^59^ and ggplot2 ^60^. Code has been deposited in GitHub: https://github.com/abcwcm/Favret2025

### ATAC-SEQ data

#### Alignment and identification of open chromatin regions

Reads were aligned to the mouse reference genome (GRCm38) with BWA-backtrack^61^. Post alignment filtering was done with samtools v1.6^62^ and Broad Institute’s Picard tools (http://broadinstitute.github.io/picard/) to remove unmapped reads, improperly paired reads, nonunique reads, and duplicates. To identify regions of open chromatin, peak calling was performed with MACS2 v2.2.6^63^. Only peaks with adjusted *P* values smaller than 0.01 were retained.

#### ATAC-SEQ peak atlas creation

Consensus peak sets were generated for tumor and infection at each transition if a peak was found in at least two replicates. The peak atlas was annotated using the ChIPseeker v1.42.1^64^ and TxDb.Mmusculus.UCSC.mm10.knownGene^65^.

#### Differentially accessible regions

Regions where the chromatin accessibility changed between different conditions were identified with DESeq2 v1.46.0, and only Benjamini–Hochberg corrected *P* values < 0.05 were considered statistically significant. A log_2_fold change cutoff of 1 was used in some analyses as indicated. When comparing earlier time points against previously published chromatin accessibility data at later time points, hidden batch effects were estimated using the svaseq function from sva v3.54.0^66^, and the top 3 surrogate variables were accounted for in DESeq2.

#### TCF1 ChIP+motif+ peak atlas creation

TCF1 ChIP data was downloaded from GSE164670^37^. Peak atlas was subset using TCF1 ChIP data selecting maximum gap distance of 500 bp between peaks to identify 17,780 TCF1 ChIP+ peaks within our peak atlas. Motif enrichment analysis was run in this subset atlas using HOMER and JASPAR2024 (DOI: <u>10.18129/B9.bioc.JASPAR2024</u>) database. Peaks containing TCF1 (JASPAR MA0769.3^41^) motif with a motif hit with log(odds ratio)>7^37^ were identified to generate a TCF1 ChIP+motif+ peak atlas of 2,419 unique peaks.

For clustered heatmap, we treated naïve as one group, pre-division samples as one group (E6h, E12h, E24h), D2 samples as one group, D5 and memory as one group. We then used our TCF1 ChIP+motif+ peak atlas and performed supervised k-Means (k=5) clustering to identify changes across these groups.

### RNA-SEQ data analysis

Adaptors were trimmed from raw sequencing reads with TrimGalore v0.6.10 (http://www.bioinformatics.babraham.ac.uk/projects/trim_galore/) and Cutadapt v2.6 ^67^. Trimmed reads were mapped with STAR v2.7.9a ^68^ to the mouse reference genome (GRCm38.p6) and fragments per gene were counted with featureCounts v2.0.1 ^69^ with respect to Gencode vM25 comprehensive gene annotations. Hidden batch effects were estimated using the svaseq function from sva v3.54.0^66^, and the top 3 surrogate variables were accounted for in DESeq2. Principial component analysis and expression heatmaps were created using batch corrected and variance-stabilizing transformed counts. LRT analysis (P<0.0001) across the different times points was used to identify genes containing the TCF1 ChIP+motif+ peaks. Heatmaps are centered and scaled by row.

## Data availability

The RNA-SEQ (GSE306744) and ATAC-SEQ (GSE306736) data have been deposited in the Gene Expression Omnibus. Previously published datasets (GSE209712)(GSE89308) were also used. All data generated and supporting the findings of this study are available within the paper or are available from the corresponding author on request.

## ACKNOWLEDGEMENTS

We thank A. Schietinger and members of the Philip laboratory for helpful discussions. We thank the Vanderbilt Division of Animal Care, D. Flaherty and the Vanderbilt University Medical Center Flow (VUMC) Cytometry Shared Resource Core, A. Jones and the Vanderbilt Technologies for Advanced Genomics (VANTAGE) Core, and Cassidy Cobbs, A. Viale and the Sloan Kettering Integrated Genomics Operation Core (IGO). We thank P. Lauer and Aduro Biotech for providing attenuated *Listeria* strains. We thank the Rathmell lab for providing human PBMCs. We thank A. Schietinger and K.M. Hawley for technical assistance with the RNA electroporation protocol. This work was supported by the following funding sources: NIH T32GM008554 (N.R.F.), V Foundation Scholar Award (M.P.), NIH R37CA263614 (M.P.), Vanderbilt-Ingram Cancer Center (VICC) SPORE Career Enhancement Program (M.P.) NIH P50CA098131, Vanderbilt Digestive Disease Research Center (VDDRC) Young Investigator and Pilot Award (M.P.) NIH P30DK058404, ASH Medical Student Physician-Scientist Award (L.A.B.), Medical Scientist Training Program (MSTP) NIH T32GM007347 to the Vanderbilt MSTP Program (M.W.R.), NIH T32CA009592 (C.R.D.R), and NIH T32AR059039 (M.M.E.). The VUMC Flow Cytometry Shared Resource is supported by the VICC (NIH P30CA68485) and the VDDRC (NIH P30DK058404). VANTAGE is supported by the VICC (NIH P30CA68485), the Vanderbilt Vision Center (NIH P30EY08126) and the NIH G20RR030956. IGO is supported by NIH P30CA08748, Cycle for Survival, and the Marie-Josée and Henry R. Kravis Center for Molecular Oncology.

## AUTHOR CONTRIBUTIONS

NRF and MP conceived and designed the study and analyzed and interpreted data. NRF carried out experiments, assisted by MMW, MWR, CNM, MME, KAM, JJR, CRDR, ZDE, LAB, and WTM. MA, PZ, and DB designed and performed computational analyses of RNA-SEQ and ATAC-SEQ data. NRF and MP wrote the manuscript, with all authors contributing to the writing and providing feedback.

## ETHICS DECLARATIONS

The authors declare no competing interests.

---

**Extended Data Figure 1.**
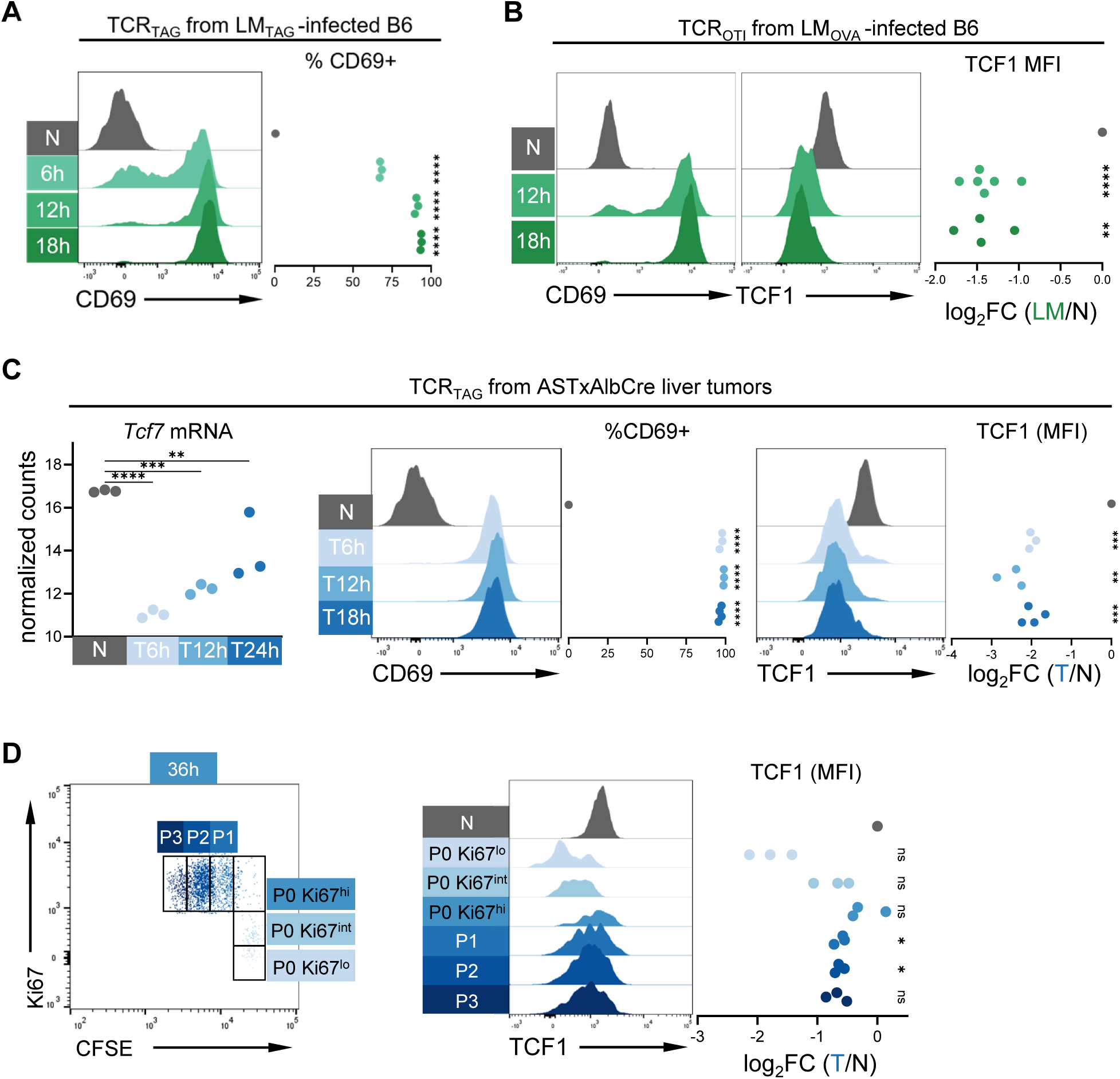
TCF1 drops upon CD8 T cell priming prior to cell division. **A.** Representative histograms and summary plot of CD69 expression on CD8^+^ Thy1.1^+^ TCR_TAG_ from spleens of infected mice (green) and naive *in vivo* control (N; grey). Each symbol represents an individual mouse with n=3 per group. \*\*\*\**P*<0.0001 determined by one sample t test. **B.** CD69 (left) and TCF1 (right) expression on CD8^+^Thy1.1^+^TCR_OTI_ analyzed 12 and 18h after transfer into LM_OVA_-infected B6 (green) and naive *in vivo* control (N; grey). Log_2_ FC of TCF1 MFI in comparison to N. \*\**P*=0.002, \*\*\*\**P*<0.0001 determined by one sample t test. **C**. CFSE-labeled naive TCR_TAG_ (Thy1.1) were adoptively transferred into ASTxAlb-Cre mice (Thy1.2) bearing late-stage liver tumors. TCR_TAG_ were re-isolated from liver tumors and analyzed by flow cytometry or sorted for sequencing at 6, 12, and 18/24 hours (h) post-transfer. Normalized counts (variance-stabilized (log_2_) expression) of *Tcf7* transcripts from TCR_TAG_ from published RNA-SEQ dataset (GSE<u>209712</u>){Rudloff, 2023 #48} (left). Representative histograms of CD69 and TCF1 expression on CD8^+^ Thy1.1^+^ TCR_TAG_ from livers of tumor-bearing mice (blue) and naive *in vivo* control (N; grey) (right). Summary plots show %CD69+ and log_2_ fold change (FC) of TCF1 MFI in comparison to N. Each symbol represents an individual mouse with n=3 per group. \*\**P*=0.0056, \*\*\**P*<0.001, \*\*\*\**P*<0.0001 determined by one sample t test. **D**. CFSE and Ki67 expression in CFSE-labeled naive TCR_TAG_ from livers of tumor-bearing mice. At 36 hours, cells are gated based on division (P0-P3) and Ki67 expression (lo, int, hi) (left). TCF1 expression of cell populations gated as shown shown (right). Data is concatenated from 3 biologic replicates per condition. Log_2_ fold change of TCF1 MFI in comparison to N. ns, not significant. \**P*<0.008 determined by one sample t test with Bonferroni correction with n=3 per group.

**Extended Data Figure 2.**
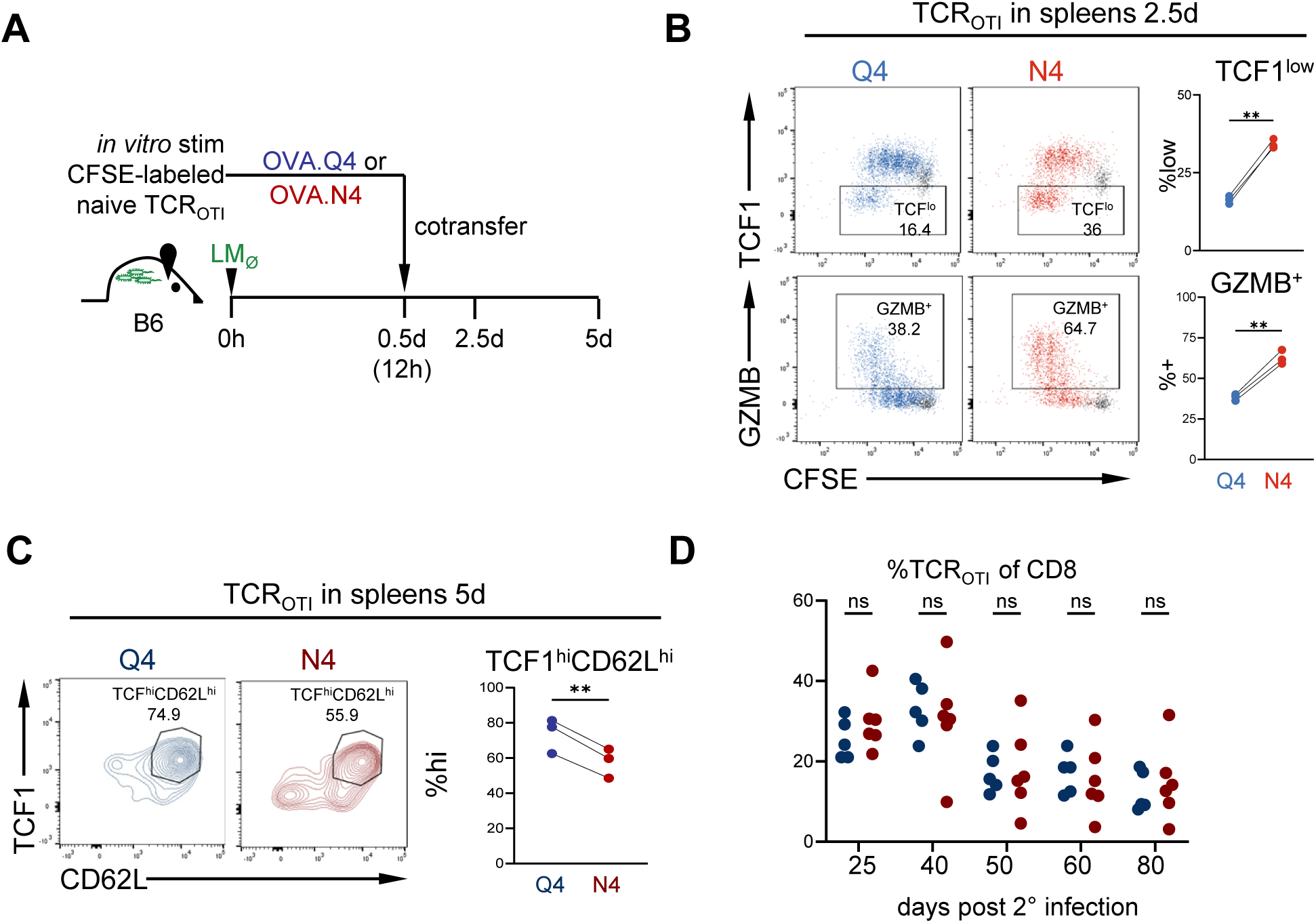
Initial T cell priming conditions impact later effector versus memory differentiation. **A.** Experimental scheme: CFSE-labeled naive TCR_OTI_ (Thy1.1 or Thy1.12) were stimulated *in vitro* with splenocytes pulsed with either low avidity OVA.Q4 (Q4;blue) or high avidity OVA.N4 (N4;red). At 12h, live CD8^+^CD90.1^+^CD69^+^ were sorted and co-adoptively transferred into LM_Ø_-infected B6 (Thy1.2). Lymphocytes were analyzed at indicated time points. **B.** CFSE by TCF1 or GZMB expression in the spleen 2.5d post-transfer, data was concatenated from 3 biologic replicates. \*\**P*<0.01 determined by paired t test. **C.** CD62L by TCF1 expression in the spleen 5d post-transfer. \*\**P*=0.0049 determined by paired t test. Data was concatenated from 3 biologic replicates. n=3 per group and representative of two independent experiments (B-C). **D.** CFSE-labeled naive TCR_OTI_ (Thy1.12) were stimulated *in vitro* with low avidity OVA.Q4 (Q4;blue) or high avidity OVA.N4 (N4;red) and 12h later, adoptively transferred into LM_OVA_-infected B6 (Thy1.2). At a memory timepoint (30+d following transfer), mice were re-challenged with LM_OVA_ (2° infection). %TCR_OTI_ of CD8 T cells in the blood at indicated days post 2° infection. NS=not significant determined by two-way ANOVA with post-hoc Sidak multiple comparison test.

**Extended Data Figure 3.**
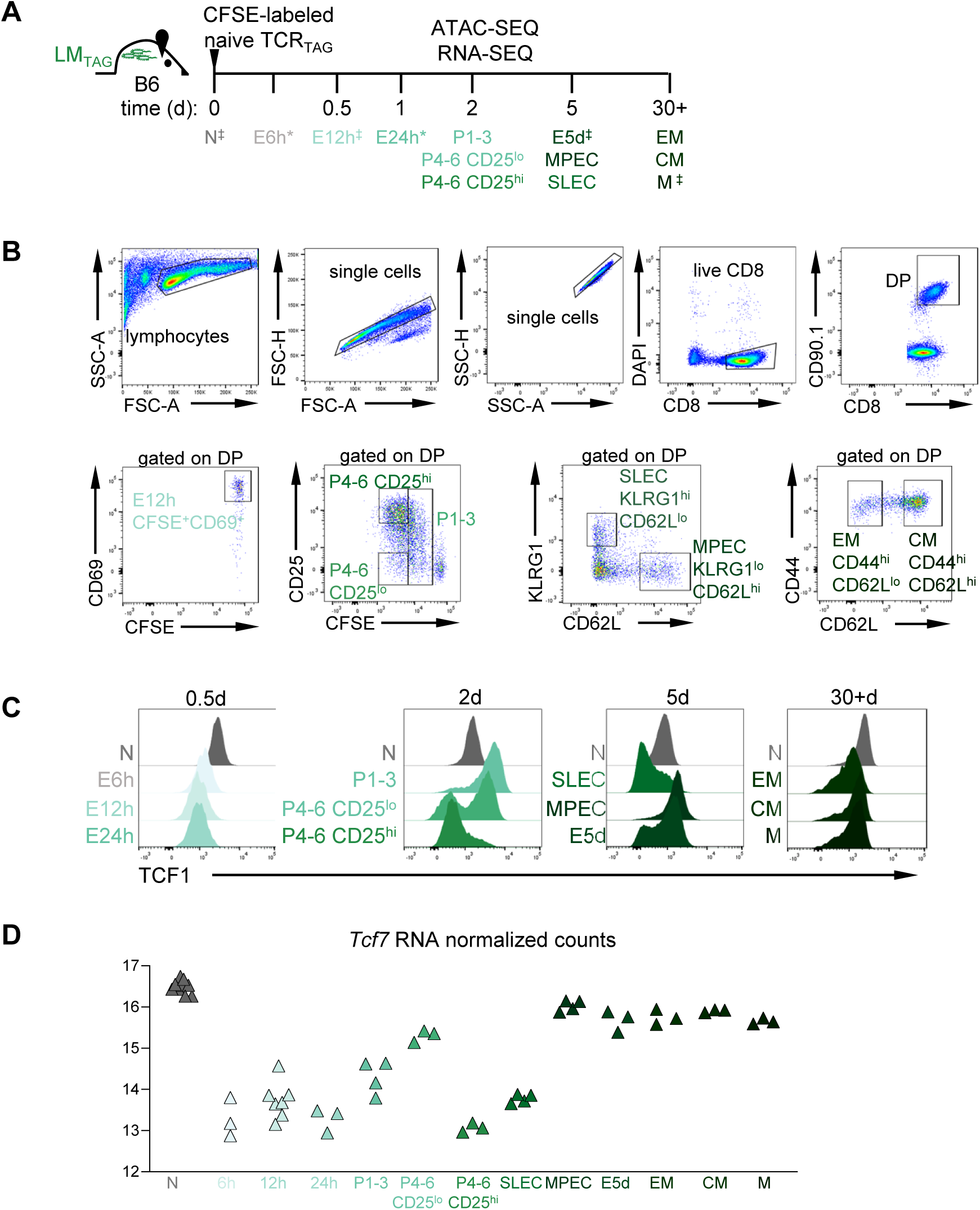
Chromatin accessibility and transcriptional analysis during CD8 T cell differentiation. **A**. (as in Fig. 5A and shown again for reference) Experimental/sorting scheme: CFSE-labeled naive TCR_TAG_(Thy1.1) were adoptively transferred into LM_TAG_-infected B6 mice; spleens were harvested at indicated time points and sorted as shown in Extended Data Fig. 3B. Time points labeled with the “*” (naive, 6h, 12h, 24h and 5d, 30+d) indicate that data was obtained from previously published studies (GSE209712)(GSE89308) ^12,38^. Time points labeled with “‡” indicate samples that were sorted in the current study and also represented in published data sets. **B**. Gating strategy to sort TCR_TAG_ from infected spleens for sequencing studies. At all timepoints, TCR_TAG_ were sorted using liveCD8^+^CD90.1^+^. At D2, samples were sorted on CFSE and CD25 expression. At D5, samples were sorted on KLRG1 and CD62L expression. At D30+, samples were sorted on CD44 and CD62L expression. **C**. TCF1 expression by flow cytometry of TCR_TAG_ at the time points and gated as shown in B. **D***. Tcf7* expression based on RNA-SEQ across time points.

**Extended Data Figure 4.**
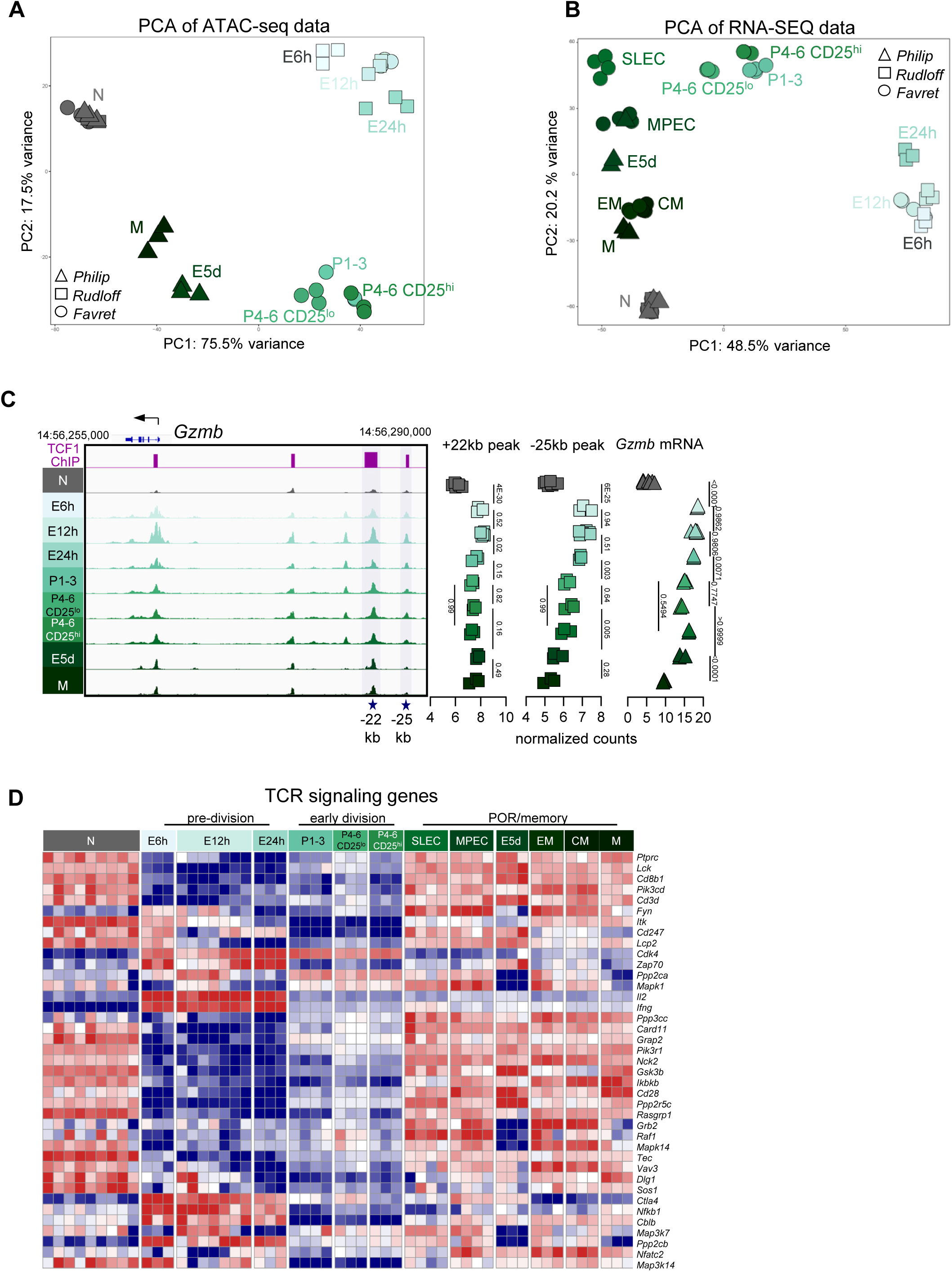
TCF1-regulated effector genes undergo priming-induced remodeling and activation while TCR signaling genes are downregulated. **A.** PCA comparing chromatin accessibility between naive and differentiated CD8 T cells by the top 1000 variable peaks. Each symbol represents a single biological replicate. Symbols indicate samples generated in the current study as well as from published studies ^12,38^(GSE209712)(GSE89308). **B.** PCA comparing gene expression changes of naive and differentiated CD8 T cells by the top 1000 variable genes. **C.** Chromatin accessibility profile across the *Gzmb* locus. Purple bars indicate TCF1 ChIP binding sites. -22kb and -25kb peaks (blue stars) and quantified (right) are TCF1 ChIP+ motif+ sites. *Gzmb* expression (right) based on RNA-SEQ across time points. *FDR* values as determined by DEseq2 shown for sequential time points. **D.** Heatmap showing TCR signaling genes (KEGG mmu04660) containing TCF1-regulated peaks that were differentially-expressed by LRT analysis (P<0.0001), z-score normalization across rows.

**Extended Data Figure 5.**
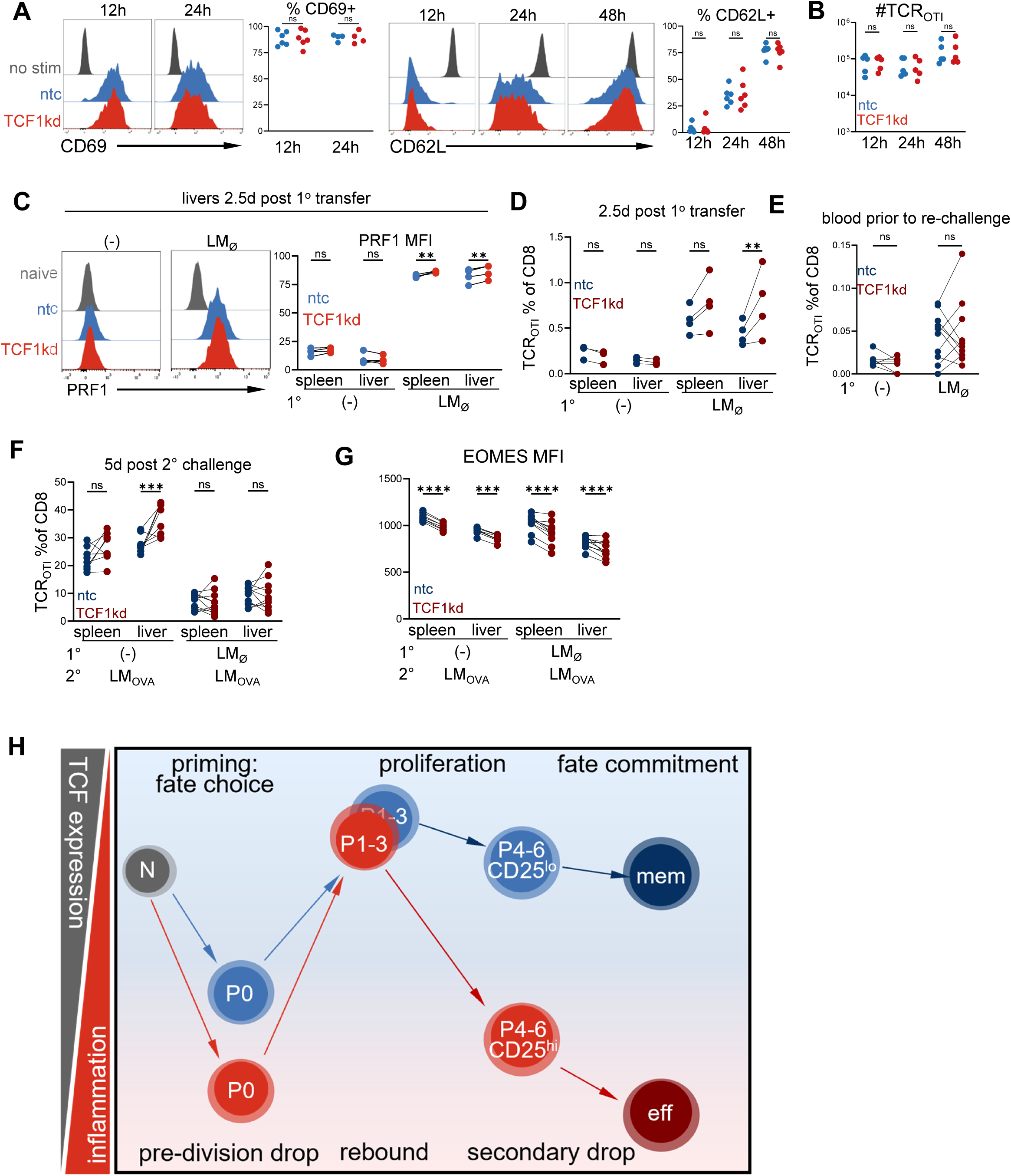
Pre-division TCF1 drop enables inflammation-driven effector differentiation. **A**. CD69 and CD62L expression on TCF1kd and ntc samples at indicated time points after Q4 stimulation as in Fig. 6A. **B**. Number of ntc and TCF1kd TCR_OTI_ *in vitro* at indicated timepoints. NS=not significant determined by two-way ANOVA with post-hoc Sidak multiple comparison test. n=4 per group, representative of two independent experiments (A,B). **C-D**. PRF1 expression on ntc and TCF1kd TCR_OTI_ (C) and %TCR_OTI_ of total CD8 T cells (D) 2.5d following transfer into B6 or LM_Ø_-infected B6 mice. \*\**P*=0.0063 determined by two-way ANOVA with post-hoc Sidak multiple comparison test. n=4 per group, representative of two independent experiments. **E**. %TCR_OTI_ of total CD8 T cells in the blood 30d post-transfer as described in Fig. 5E. Statistical analysis by two-way ANOVA with post-hoc Sidak multiple comparison test. **F**. %TCR_OTI_ of total CD8 T cells in the spleen and liver 5d following 2° challenge with LM_OVA_. \*\*\**P*= 0.0003 determined by two-way ANOVA with post-hoc Sidak multiple comparison test. **G**. EOMES expression 5d 2° challenge with LM_OVA_. \*\*\**P*<0.001, \*\*\*\**P*<0.0001 determined by two-way ANOVA with post-hoc Sidak multiple comparison test. n=8-10, representative of two independent experiments. **H**. Proposed model: Upon priming of naive CD8 T cells and prior to cell division (P0), TCF1 expression drops, with high TCR signal strength and/or inflammatory signaling causing a large drop (red), and low TCR signal strength and inflammatory signaling causing a more modest drop (blue). The TCF1 drop causes chromatin remodeling in TCF1-regulated genes, poising the TCF1-dropped T cells to express effector genes and inflammatory cytokine receptors. Upon cell cycle entry (P1-3), all T cells undergo a high rebound of TCF1 expression. With proliferation (P4-6), T cells which underwent a larger pre-division TCF1 drop will be more sensitive to inflammatory cytokines, undergoing a secondary TCF1 drop and commitment to the effector fate (dark red). T cells which underwent a minimal TCF1 drop are less responsive to inflammatory cytokines and do not undergo a secondary TCF1 drop, instead committing to the memory fate (dark blue).

## REFERENCES

1 Gerlach, C. et al. Heterogeneous differentiation patterns of individual CD8+ T cells. Science 340, 635–639 (2013).

2 Buchholz, V. R. et al. Disparate individual fates compose robust CD8+ T cell immunity. Science 340, 630–635 (2013).

3 Ahmed, R., Bevan, M. J., Reiner, S. L. & Fearon, D. T. The precursors of memory: models and controversies. Nat Rev Immunol 9, 662–668 (2009).

4 Chang, J. T. et al. Asymmetric T lymphocyte division in the initiation of adaptive immune responses. Science 315, 1687–1691 (2007).

5 Chang, J. T. et al. Asymmetric proteasome segregation as a mechanism for unequal partitioning of the transcription factor T-bet during T lymphocyte division. Immunity 34, 492–504 (2011).

6 Rudloff, M. W. et al. Hallmarks of CD8(+) T cell dysfunction are established within hours of tumor antigen encounter before cell division. Nat Immunol 24, 1527–1539 (2023).

7 Stadhouders, R., Filion, G. J. & Graf, T. Transcription factors and 3D genome conformation in cell-fate decisions. Nature 569, 345–354 (2019).

8 Kaech, S. M. & Cui, W. Transcriptional control of effector and memory CD8+ T cell differentiation. Nat Rev Immunol 12, 749–761 (2012).

9 Escobar, G., Mangani, D. & Anderson, A. C. T cell factor 1: A master regulator of the T cell response in disease. Sci Immunol 5 (2020).

10 Gounari, F. & Khazaie, K. TCF-1: a maverick in T cell development and function. Nat Immunol 23, 671–678 (2022).

11 Verbeek, S. et al. An HMG-box-containing T-cell factor required for thymocyte differentiation. Nature 374, 70–74 (1995).

12 Willinger, T. et al. Human naive CD8 T cells down-regulate expression of the WNT pathway transcription factors lymphoid enhancer binding factor 1 and transcription factor 7 (T cell factor-1) following antigen encounter in vitro and in vivo. Journal of immunology 176, 1439–1446 (2006).

13 Gattinoni, L. et al. Wnt signaling arrests effector T cell differentiation and generates CD8+ memory stem cells. Nat Med 15, 808–813 (2009).

14 Scharer, C. D., Barwick, B. G., Youngblood, B. A., Ahmed, R. & Boss, J. M. Global DNA methylation remodeling accompanies CD8 T cell effector function. Journal of immunology 191, 3419–3429 (2013).

15 Tiemessen, M. M. et al. T Cell factor 1 represses CD8+ effector T cell formation and function. Journal of immunology 193, 5480–5487 (2014).

16 Lin, W. W. et al. CD8+ T Lymphocyte Self-Renewal during Effector Cell Determination. Cell reports 17, 1773–1782 (2016).

17 Staveley-O’Carroll, K. et al. In vivo ligation of CD40 enhances priming against the endogenous tumor antigen and promotes CD8+ T cell effector function in SV40 T antigen transgenic mice. Journal of immunology 171, 697–707 (2003).

18 Lin, W. W. et al. CD8(+) T Lymphocyte Self-Renewal during Effector Cell Determination. Cell Rep 17, 1773–1782 (2016).

19 Danilo, M., Chennupati, V., Silva, J. G., Siegert, S. & Held, W. Suppression of Tcf1 by Inflammatory Cytokines Facilitates Effector CD8 T Cell Differentiation. Cell reports 22, 2107–2117 (2018).

20 Simonetti, S. et al. A DNA/Ki67-Based Flow Cytometry Assay for Cell Cycle Analysis of Antigen-Specific CD8 T Cells in Vaccinated Mice. J Vis Exp (2021).

21 Miller, I. et al. Ki67 is a Graded Rather than a Binary Marker of Proliferation versus Quiescence. Cell reports 24, 1105–1112 e1105 (2018).

22 Wolf, T. et al. Dynamics in protein translation sustaining T cell preparedness. Nat Immunol 21, 927–937 (2020).

23 Wu, T., et al. The TCF1-Bcl6 axis counteracts type I interferon to repress exhaustion and maintain T cell stemness. Sci Immunol 1 (2016).

24 Silva, J. G., et al. Emergence and fate of stem cell-like Tcf7(+) CD8(+) T cells during a primary immune response to viral infection. Sci Immunol 8, eadh3113 (2023).

25 Shakiba, M. et al. TCR signal strength defines distinct mechanisms of T cell dysfunction and cancer evasion. J Exp Med 219 (2022).

26 Rosette, C. et al. The impact of duration versus extent of TCR occupancy on T cell activation: a revision of the kinetic proofreading model. Immunity 15, 59–70 (2001).

27 Hsieh, C. S. et al. Development of TH1 CD4+ T cells through IL-12 produced by Listeria-induced macrophages. Science 260, 547–549 (1993).

28 Mescher, M. F. et al. Signals required for programming effector and memory development by CD8+ T cells. Immunol Rev 211, 81–92 (2006).

29 Zehn, D., Lee, S. Y. & Bevan, M. J. Complete but curtailed T-cell response to very low-affinity antigen. Nature 458, 211–214 (2009).

30 Joshi, N. S. et al. Inflammation directs memory precursor and short-lived effector CD8(+) T cell fates via the graded expression of T-bet transcription factor. Immunity 27, 281–295 (2007).

31 Zangari, B. et al. Tcf-1 protects anti-tumor TCR-engineered CD8(+) T-cells from GzmB mediated self-destruction. Cancer Immunol Immunother 71, 2881–2898 (2022).

32 Johnnidis, J. B., et al. Inhibitory signaling sustains a distinct early memory CD8(+) T cell precursor that is resistant to DNA damage. Sci Immunol 6 (2021).

33 Johnson, J. L. et al. Lineage-Determining Transcription Factor TCF-1 Initiates the Epigenetic Identity of T Cells. Immunity 48, 243–257 e210 (2018).

34 Shan, Q. et al. Tcf1-CTCF cooperativity shapes genomic architecture to promote CD8(+) T cell homeostasis. Nat Immunol 23, 1222–1235 (2022).

35 Wang, W. et al. TCF-1 promotes chromatin interactions across topologically associating domains in T cell progenitors. Nat Immunol 23, 1052–1062 (2022).

36 Xing, S. et al. Tcf1 and Lef1 transcription factors establish CD8(+) T cell identity through intrinsic HDAC activity. Nat Immunol 17, 695–703 (2016).

37 Shan, Q. et al. Tcf1 and Lef1 provide constant supervision to mature CD8(+) T cell identity and function by organizing genomic architecture. Nature communications 12, 5863 (2021).

38 Buenrostro, J. D., Giresi, P. G., Zaba, L. C., Chang, H. Y. & Greenleaf, W. J. Transposition of native chromatin for fast and sensitive epigenomic profiling of open chromatin, DNA-binding proteins and nucleosome position. Nature methods 10, 1213–1218 (2013).

39 Kalia, V. et al. Prolonged interleukin-2Ralpha expression on virus-specific CD8+ T cells favors terminal-effector differentiation in vivo. Immunity 32, 91–103 (2010).

40 Philip, M. et al. Chromatin states define tumour-specific T cell dysfunction and reprogramming. Nature 545, 452–456 (2017).

41 Isakova, A. et al. SMiLE-seq identifies binding motifs of single and dimeric transcription factors. Nature methods 14, 316–322 (2017).

42 Man, K. et al. Transcription Factor IRF4 Promotes CD8(+) T Cell Exhaustion and Limits the Development of Memory-like T Cells during Chronic Infection. Immunity 47, 1129–1141 e1125 (2017).

43 Harly, C. et al. A Shared Regulatory Element Controls the Initiation of Tcf7 Expression During Early T Cell and Innate Lymphoid Cell Developments. Frontiers in immunology 11, 470 (2020).

44 Weber, B. N. et al. A critical role for TCF-1 in T-lineage specification and differentiation. Nature 476, 63–68 (2011).

45 Evangelista, J. E. et al. Enrichr-KG: bridging enrichment analysis across multiple libraries. Nucleic Acids Res 51, W168–W179 (2023).

46 Kallies, A., Xin, A., Belz, G. T. & Nutt, S. L. Blimp-1 transcription factor is required for the differentiation of effector CD8(+) T cells and memory responses. Immunity 31, 283–295 (2009).

47 Dominguez, C. X. et al. The transcription factors ZEB2 and T-bet cooperate to program cytotoxic T cell terminal differentiation in response to LCMV viral infection. J Exp Med 212, 2041–2056 (2015).

48 Youngblood, B. et al. Effector CD8 T cells dedifferentiate into long-lived memory cells. Nature 552, 404–409 (2017).

49 Jeannet, G. et al. Essential role of the Wnt pathway effector Tcf-1 for the establishment of functional CD8 T cell memory. Proc Natl Acad Sci U S A 107, 9777–9782 (2010).

50 Zhou, X. et al. Differentiation and persistence of memory CD8(+) T cells depend on T cell factor 1. Immunity 33, 229–240 (2010).

51 Shan, Q. et al. Tcf1 preprograms the mobilization of glycolysis in central memory CD8(+) T cells during recall responses. Nat Immunol 23, 386–398 (2022).

52 Abadie, K. et al. Reversible, tunable epigenetic silencing of TCF1 generates flexibility in the T cell memory decision. Immunity 57, 271–286 e213 (2024).

53 Hogquist, K. A. et al. T cell receptor antagonist peptides induce positive selection. Cell 76, 17–27 (1994).

54 Schietinger, A. et al. Tumor-Specific T Cell Dysfunction Is a Dynamic Antigen-Driven Differentiation Program Initiated Early during Tumorigenesis. Immunity 45, 389–401 (2016).

55 Stahl, S. et al. Tumor agonist peptides break tolerance and elicit effective CTL responses in an inducible mouse model of hepatocellular carcinoma. Immunol Lett 123, 31–37 (2009).

56 Brockstedt, D. G. et al. Listeria-based cancer vaccines that segregate immunogenicity from toxicity. Proc Natl Acad Sci U S A 101, 13832–13837 (2004).

57 FastQC: A Quality Control Tool for High Throughput Sequence Data (2010).

58 Hartley, S. W. & Mullikin, J. C. QoRTs: a comprehensive toolset for quality control and data processing of RNA-Seq experiments. BMC Bioinformatics 16, 224 (2015).

59 R: A Language and Environment for Statistical Computing (R Foundation for Statistical Computing, 2023).

60 Wickham, H. ggplot2: Elegant Graphics for Data Analysis. (2016).

61 Li, H. & Durbin, R. Fast and accurate short read alignment with Burrows-Wheeler transform. Bioinformatics 25, 1754–1760 (2009).

62 Li, H. et al. The Sequence Alignment/Map format and SAMtools. Bioinformatics 25, 2078–2079 (2009).

63 Liu, T. Use model-based Analysis of ChIP-Seq (MACS) to analyze short reads generated by sequencing protein-DNA interactions in embryonic stem cells. Methods in molecular biology 1150, 81–95 (2014).

64 Yu, G., Wang, L. G. & He, Q. Y. ChIPseeker: an R/Bioconductor package for ChIP peak annotation, comparison and visualization. Bioinformatics 31, 2382–2383 (2015).

65 TxDb.Mmusculus.UCSC.mm10.knownGene: Annotation package for TxDb object(s). (R package version 3.4.7., 2019).

66 sva: Surrogate Variable Analysis. (R package version 3.44.0, 2022).

67 Martin, M. Cutadapt removes adapter sequences from high-throughput sequencing reads. EMBnet.journal 17 (2011).

68 Dobin, A. et al. STAR: ultrafast universal RNA-seq aligner. Bioinformatics 29, 15–21 (2013).

69 Liao, Y., Smyth, G. K. & Shi, W. featureCounts: an efficient general purpose program for assigning sequence reads to genomic features. Bioinformatics 30, 923–930 (2014).

